# First in, first folded: evolution of vectorial substrate folding by metazoan Hsp60/10

**DOI:** 10.64898/2026.11.23.768786

**Authors:** Manisha Kochar, Shikha Rao, Sarbani Bhattacharjee, Purva Goel, Himanshu Timsina, Md Khabeer Azhar, Vishal Bharti, Arjun Ray, Dilip Kumar, Koyeli Mapa, Kausik Chakraborty

**Affiliations:** CSIR-Institute of Genomics and Integrative Biology, New Delhi 110025, India; Academy of Scientific and Innovative Research (AcSIR), Ghaziabad 201002, India; Department of Life Sciences, School of Natural Sciences, Shiv Nadar Institution of Eminence, Delhi-NCR, Greater Noida, Uttar Pradesh, 201314, India; Bhaskaracharya College of Applied Sciences, University of Delhi, New Delhi 110075, India; Department of Biotechnology, Jamia Millia Islamia, New Delhi 110025, India; Department of Computational Biology, Indraprastha Institute of Information Technology-Delhi, New Delhi 110020, India; Trivedi School of Biosciences, Ashoka University, Sonipat, Haryana 131029, India

**Keywords:** chaperonin, Hsp60, GroEL, protein folding, vectorial folding, co-translational folding, mitochondrial protein import, proteostasis, molecular evolution, encapsulation

## Abstract

Group-I chaperonins such as GroEL/GroES fold proteins by encapsulating a complete non-native chain within a central chamber. This limits its substrates to proteins that can be accommodated within the cavity. Whether human mitochondrial Hsp60/10, a descendant of bacterial GroEL/ES, serving a larger proteome, retains the size constraint or bypasses it has remained unknown. Here, we show that human Hsp60/10 assists the folding of large multidomain proteins without the size-dependent decline in folding efficiency as observed with bacterial GroEL/ES. The human Hsp60/10 chaperonin assists the folding of large substrates not by expanding its cavity, but by preferentially folding the N-terminal domain of a substrate. This positional preference is absent during the refolding of fully synthesised proteins but observed only when folding is coupled to synthesis. This preference is maintained *in vivo* within the mitochondrial matrix, where the N-termini of precursor proteins are imported first. Comparative analysis of a wide array of chaperonin homologues further reveals that this mode of size-independent, N-terminally directed folding is confined to the metazoan lineage. Our findings identify vectorial folding as an evolutionary adaptation of the ancestral chaperonin system, enabling metazoan Hsp60/10s to assist the folding of larger, multidomain proteins.

**HIGHLIGHTS:**

- Human Hsp60/10 assists folding of multi-domain proteins beyond the substrate-size constraints
- Human Hsp60/10 preferentially assists the folding of N-terminal domains in multi-domain proteins
- Size-independent directional substrate folding assistance emerged in the metazoan lineage

## INTRODUCTION

The Hsp60/Hsp10 family of group-I chaperonins is an essential, ATP-dependent folding system, conserved across almost all forms of life (1–6). The bacterial GroEL/GroES from *E. coli* has served as the canonical model for chaperonin function (4, 7). During each ATP-driven cycle, GroEL captures a non-native polypeptide chain, encapsulates it in the cavity upon GroES binding, and releases it after a brief folding interval (8–11). This mechanism imposes a geometric constraint on substrate size, limiting efficient encapsulation to proteins of ∼55-60 kDa (12, 13). Consistent with this size constraint, the obligate GroEL substrates in *E. coli* cluster below this threshold (14, 15). Larger substrates have been reported (16, 17), though they remain uncommon exceptions given the distribution of substrate sizes.

Whether this size constraint applies equally to eukaryotic chaperonins remains unclear. Group I chaperonins, including GroEL, can engage substrates co-translationally (11, 18). However, engagement of an emerging polypeptide chain does not necessarily overcome the size limit imposed by encapsulation; the obligate substrates of GroEL still cluster below this size (15). Also, the reported ability of GroEL to assist the folding of larger substrates through multiple cycles of binding and release in trans remains an isolated exception rather than an established general mechanism (16). The eukaryotic cytosolic chaperonin TRiC, by contrast, folds proteins larger than its chamber by enclosing individual domains rather than encapsulating the entire polypeptide chain (19). However, TRiC is a Group II chaperonin, which differs fundamentally from Group I chaperonins in its architecture. Its hetero-oligomeric ring and built-in lid facilitate domain-wise substrate recognition and folding. Eukaryotes further exploit co-translational, domain-wise folding that is mechanistically different from the bacterial chaperonins (20, 21).

This raises an important question about the evolution of group-I chaperonins. Intriguingly, human mitochondrial Hsp60/10 descends from the bacterial GroEL/ES system, yet it serves a proteome of considerably larger, multidomain proteins. Most mitochondrial matrix proteins are synthesised in the cytosol as precursors and imported through the TIM23 translocon in an N-terminus-first direction (22–24). The imported precursor proteins are presented to the chaperonin vectorially as they enter the matrix. Does the size constraint imposed by encapsulation represent an intrinsic feature of the group-I chaperonin mechanism, shared across all its homologues, or have some lineages evolved alternative modes of substrate folding that overcome this constraint?

Here, we show that human Hsp60/10 (hHsp60/10) assists the folding of substrates beyond the size limit of its bacterial ancestor, not by expanding its central cavity, but by altering how it engages and folds its substrates. Rather than requiring encapsulation of an entire multidomain protein, Hsp60/10 preferentially assists the folding of the N-terminal domain, as the polypeptide chain emerges. Using folding reporters in *E. coli*, chaperone-free reconstituted translation, folding assays in human mitochondria, and a panel of chaperonin homologues, we find that this positional preference is absent during refolding of complete proteins but emerges when folding is coupled to protein synthesis. This mode of folding is also retained in the mitochondrial matrix, where precursor proteins are imported with an N-terminus first direction. Comparative analysis of chaperonin homologues spanning the tree of life further reveals that this size-independent, N-terminally directed folding is restricted to the metazoan lineage. Together, these findings identify the positional, N-terminally directed folding as an evolutionary innovation of the metazoan group-I chaperonins.

## RESULTS

### Human Hsp60/10 efficiently assists folding of multi-domain proteins *in vivo*

Productive folding by the conserved group-I chaperonins is thought to depend on complete encapsulation of their substrates within the chaperonin cavity (9, 13, 25–27). This requirement imposes a geometric constraint on the substrate size, with efficient encapsulation by *E. coli* GroEL/ES generally limited to proteins of ∼55-60 kDa (12, 13). Consistent with this constraint, obligate GroEL substrates of *E. coli* predominantly fall below this threshold (14, 15). The mitochondrial proteome, however, differs substantially from the bacterial cytosol, and human mitochondrial Hsp60/10, despite its evolutionary relationship with bacterial GroEL/ES, operates in a proteome containing considerably larger proteins (Figure 1A, B, Table S1). To understand whether this difference is reflected in their substrate repertoires, we compared the reported interactomes of GroEL and hHsp60 (15, 28). While the GroEL interactome showed a sharp decline in proteins larger than ∼60 kDa, hHsp60 interactors extended well beyond 80 kDa, with ∼22% exceeding the canonical encapsulation limit (Figure 1C, Table S1) (15, 28). This difference raised the possibility that hHsp60/10 has evolved to accommodate larger substrates through a mechanism distinct from that of its bacterial counterpart.

**Figure 1.**
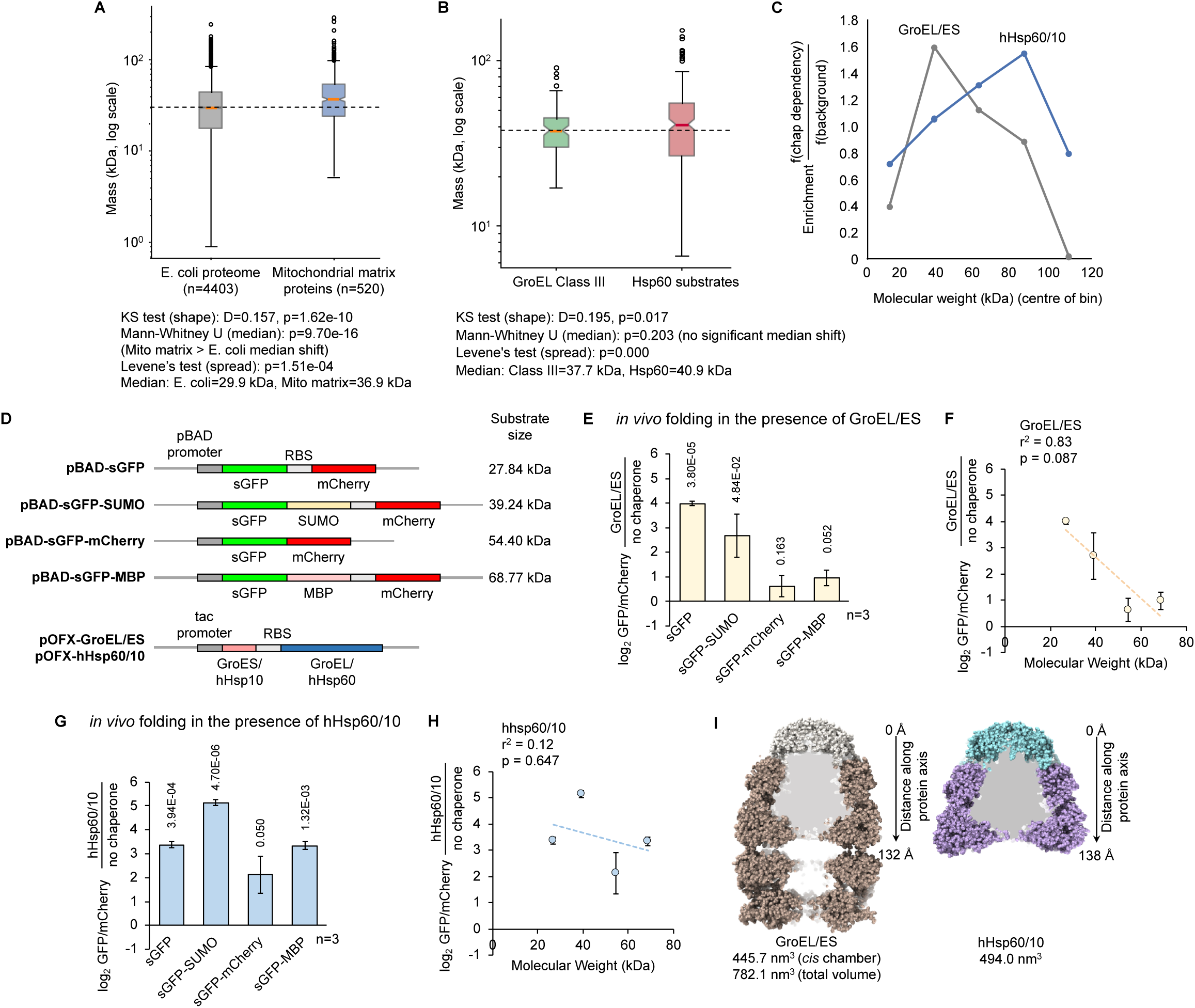
hHsp60/10 maintains folding assistance towards large multidomain substrates. **(A**) Comparison of length between curated human mitochondrial matrix targeted proteins and *E. coli* proteome after removal of mitochondrial targeting sequences. **(B)** Enrichment of substrate proteins across molecular-weight bins relative to the corresponding background proteomes. The last bin includes proteins ≥ 100kDa. **(C)** Comparison of obligate GroEL substrates and high-incidence hHsp60 interactors. Although the median substrate size is similar, the Hsp60 interactome shows significantly greater variance, consistent with an expanded substrate range. **(D)** Schematic representation of substrate and chaperone constructs used for *in vivo* folding. **(E)** Folding of fusion proteins by GroEL/ES measured *in vivo* using the GFP/mCherry reporter constructs by flow cytometry. **(F)** Correlation between GroEL/ES-assisted folding and substrate molecular weight. **(G)** Folding of the fusion proteins by hHsp60/10 *in vivo*. **(H)** Correlation between hHsp60/10-assisted folding and substrate molecular weights. Unlike GroEL/ES, hHsp60/10 activity does not correlate with substrate size. **(I)** ATP-bound structures of GroEL/ES (PDB:8BM1) and hHsp60/10 (PDB:9ES1) used for cavity analysis. Cavity volumes were calculated for the GroEL/ES double-ring complex, the GroEL/ES cis chamber alone, and the single-ring hHsp60/10 complex. The “distance along protein axis” used in Figure S1D is denoted along the structures. Data are mean ± SD from three independent experiments. Statistical significance was determined using two-tailed Student’s t-tests; p-values relative to the respective spontaneous folding in (E, G) are shown above each bar; correlations were assessed by linear regression.

To understand how substrate size influences chaperonin-assisted folding, we developed a series of multidomain substrates using slow-folding GFP (sGFP; yeGFP K45E) as a common folding reporter (29). sGFP is a destabilised GFP variant that folds inefficiently on its own but can be efficiently assisted by GroEL/ES, providing a well-characterised system for quantifying chaperonin-dependent folding *in vivo* and *in vitro* (Figure S1A) (29, 30). We fused sGFP to partner domains of increasing molecular size (SUMO, mCherry or MBP) at its C-terminus, thereby varying the overall substrate size while retaining the same folding-defective reporter domain (Figure 1D)

Because productive folding of sGFP is coupled to chromophore maturation, GFP fluorescence provides a quantitative readout of native-state acquisition. Previous work established that GroEL/ES actively promotes sGFP folding rather than merely preventing its aggregation (29, 30). To quantify folding in vivo, sGFP constructs were expressed from an operon with mCherry translated from the same mRNA, which served as an internal control for transcriptional and translational variation (Figure 1D). The resulting GFP/mCherry ratio therefore reports the folding efficiency of sGFP, and comparing this ratio in the absence and presence of an overexpressed chaperonin quantifies the assistance provided by that chaperonin (Figure 1D, S1B). For constructs in which mCherry is fused to sGFP, mCherry fluorescence did not serve as an internal control as sGFP folding would determine the in vivo stability of the fusion protein and hence also the mCherry fluorescence.

Consistent with previous reports, GroEL/ES efficiently enhanced folding of sGFP in wild-type *E. coli* (K12 strain BW25113; hereafter Wtk) (Figure 1E). However, all fusion constructs were assisted less efficiently than the single-domain sGFP, indicating reduced enhancement of sGFP folding within a multi-domain construct (Figure 1E). The folding assistance showed a negative trend with increasing molecular size of the fusion proteins (Figure 1F), consistent with earlier observations that increasing substrate size compromises productive folding by GroEL/ES (12, 13, 31).

In contrast, hHsp60/10 exhibited a different pattern of substrate assistance. Although hHsp60/10 enhanced sGFP folding in all multidomain contexts (Figure 1G), unlike GroEL/ES, no overall relationship between folding assistance and substrate size was observed (Figure 1H). This activity depended on the co-chaperonin, as removal of hHsp10 abolished the observed enhancement in folding (Figure S1C).

We next considered whether the ability of hHsp60/10 to assist larger substrates could be explained by an expanded folding cavity. Volumetric analysis of the published structures showed that the ATP/Hsp10-bound cis chamber of hHsp60/10 is comparable in volume to that of GroEL/ES (Figure 1I, S1D). Although the single-ring mutant of GroEL (SR1) forms a larger cis chamber than double-ring GroEL/ES (32), the predominantly single-ring assembly of hHsp60 does not result in a larger folding chamber. Hence, the ability of hHsp60/10 to assist larger substrates cannot be explained by an expanded folding cavity.

These observations point towards a functional divergence between the two chaperonin systems. While GroEL/ES assistance tends to decline as substrate size increases, hHsp60/10 can efficiently assist folding within multidomain proteins without a corresponding dependence on total molecular size.

### N-terminal domains are preferred substrates of hHsp60 irrespective of total size

We next tested whether folding assistance is influenced by the position of the folding-defective domain within the polypeptide. To assess these variables, we expanded our substrate panel of fusion proteins in which sGFP was appended to partner proteins (SUMO, mCherry or MBP) at either the N- or the C-terminus (Figure 2A). All constructs were designed in such a way that their total molecular weights remained within the size range compatible with encapsulation by the GroEL/ES chamber (3), allowing us to evaluate the influence of domain order and of increasing substrate size without confounding by encapsulation capacity.

**Figure 2.**
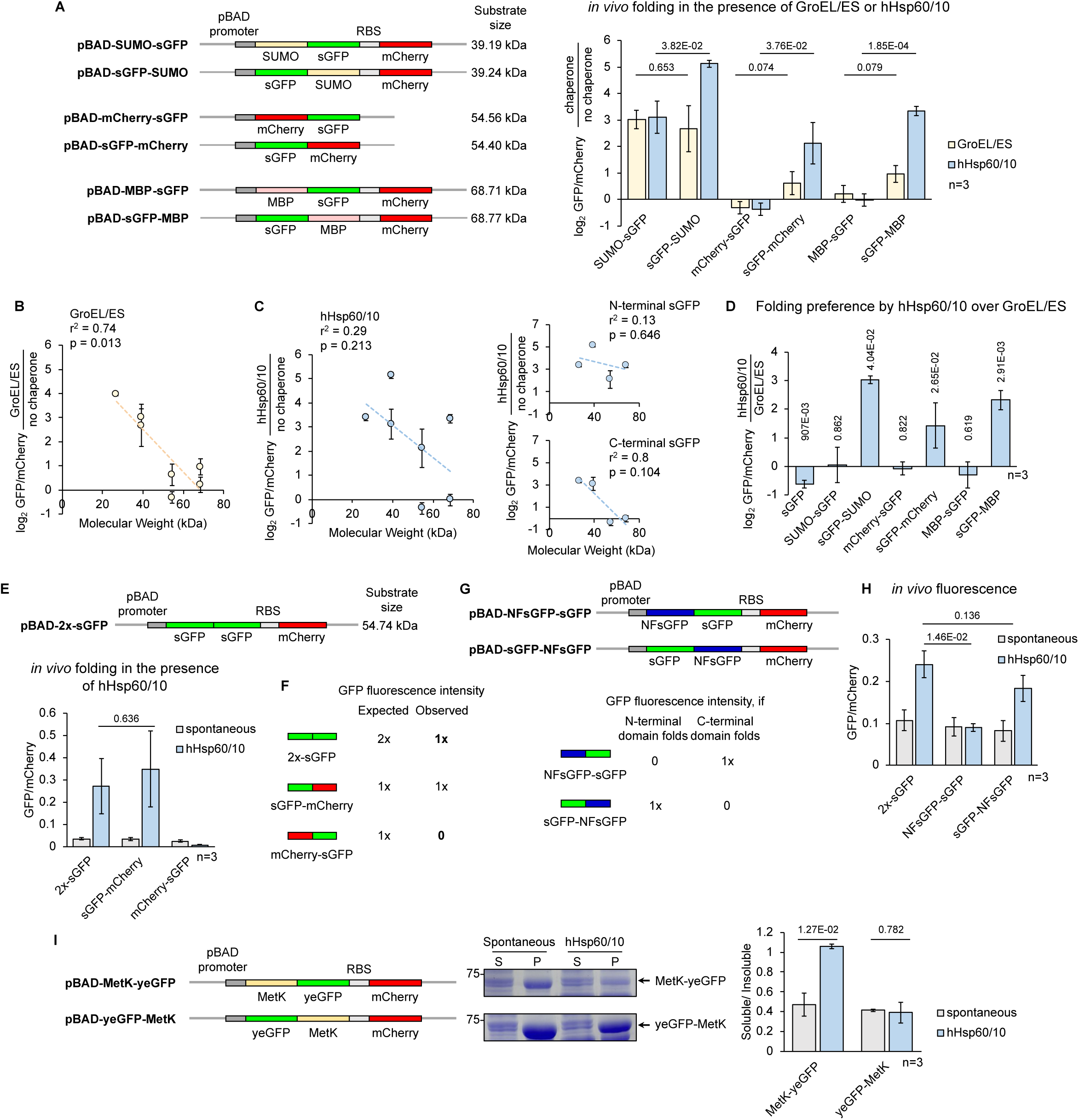
Domain order determines folding assistance by hHsp60/10. **(A**) Left, schematic representation of different fusion proteins and their predicted molecular masses. Right, folding of fusion proteins by GroEL/ES or hHsp60/10 measured *in vivo* using the GFP/mCherry reporter constructs by flow cytometry. **(B)** Correlation between GroEL/ES-assisted folding and substrate molecular weight. **(C)** Left, correlation between hHsp60/10-assisted folding and substrate molecular weights. Right, relationship between hHsp60/10 assistance and substrate size when sGFP is present at the N-terminus (top) or C-terminus (bottom) of the fusion protein. **(D)** Ratio of hHsp60/10 assisted folding to GroEL/ES assisted folding for each substrate, highlighting the relative folding preference. **(E)** Folding of tandem sGFP, sGFP-mCherry^#^ and mCherry-sGFP^#^ constructs by hHsp60/10. **(F)** Schematic showing the expected GFP fluorescence if one or both sGFP domains fold within tandem fusion proteins. **(G)** Experimental strategy using tandem fluorescent and non-fluorescent sGFP (NFsGFP) variants to distinguish preferential folding of the N-or C-terminal sGFP domain. **(H)** Folding of tandem sGFP reporters by hHsp60/10. Median GFP/mCherry fluorescence demonstrates preferential folding of the N-terminal sGFP domain by hHsp60/10. **(I)** Solubility analysis of MetK-yeGFP and yeGFP-MetK following expression in the absence or presence of hHsp60/10. Soluble (S) and insoluble (P) fractions were analysed by SDS-PAGE and corresponding band intensities were quantified (right). Data are mean ± SD from three independent experiments. Statistical significance was determined using two-tailed Student’s t-tests; p-values are shown above each bar; correlations were assessed using linear regression. ^#^Independently translating mCherry was absent from constructs where mCherry was used as a fusion partner.

For GroEL/ES, there was a size-dependent decrease in the assistance, indicating reduced enhancement of sGFP folding within a multidomain context (Figure 2A). Across the expanded substrate panel, assistance showed a negative trend with increasing molecular size (Figure 2A, B). SUMO-sGFP, which was assisted less than sGFP by GroEL/ES (Figure 1E), was efficiently encapsulated by GroEL/ES in the presence of ATP·AlFx, as indicated by the expected increase in fluorescence anisotropy upon encapsulation (Figure S2A). Thus, complete failure of encapsulation of substrate did not dictate the decreasing assistance of GroEL/ES to larger substrates. Additionally, because all constructs remained within the encapsulation-compatible size range, this decline cannot be attributed to a complete failure of encapsulation, although the efficiency of encapsulation may be different. These observations suggest that increasing substrate size progressively lowers the probability of productive folding within the GroEL/ES cavity.

hHsp60/10, however, showed a striking dependence on domain orientation (Figure 2A). Although no overall relationship between folding assistance and substrate size was observed, a clear trend emerged when the entire dataset was analysed collectively, with constructs grouped according to the position of sGFP within the fusion protein (Figure 2C). Assistance provided by hHsp60/10 reduced progressively with size when sGFP occupied the C-terminal position (SUMO-sGFP, mCherry-sGFP or MBP-sGFP) (Figure 2C). In contrast, robust enhancement in folding assistance was observed when sGFP was positioned at the N-terminus (sGFP-SUMO, sGFP-mCherry or sGFP-MBP), irrespective of the total size of the fusion protein (Figure 2C). Even constructs containing large C-terminal partner domains were assisted as efficiently as smaller substrates. Direct comparison of the two chaperonin systems further highlighted this distinction. While GroEL/ES was more effective towards isolated sGFP, hHsp60/10 was better than GroEL/ES at folding sGFP when it occupied the N-terminal position in a multidomain construct (Figure 2D).

These findings indicated that domain position, rather than total molecular size alone, strongly influences hHsp60/10-mediated folding. Whereas GroEL/ES showed a general decline in assistance with increasing substrate size, regardless of domain orientation, hHsp60/10 preferentially assisted N-terminal sGFP even when fused to substantially larger partner domains. This positional asymmetry suggested that the human mitochondrial chaperonin may engage multidomain substrates in a directional manner.

In all the constructs described above, the partner domains differed not only in size but also in their intrinsic folding properties, raising the possibility that these differences contributed to the observed positional bias. To determine whether domain order alone was sufficient to influence folding, we generated a symmetric tandem construct consisting of two identical sGFP domains fused in series (2x-sGFP) (Figure 2E). This design eliminated differences in domain identity, size and stability between the two positions. If domain position does not influence hHsp60/10 activity, both sGFP domains would be expected to fold with similar probability. Conversely, if assistance is directional, fluorescence would predominantly reflect folding of a single domain.

Co-expression of 2x-sGFP with hHsp60/10 increased the fluorescence well above the spontaneous level, confirming that the tandem substrate remained competent for chaperonin-assisted folding (Figure 2E). However, the magnitude of this signal corresponded to that expected from a single folded sGFP domain rather than two independently folded domains (Figure 2E, F). The fluorescence from 2x-sGFP was comparable to that of the similarly sized sGFP-mCherry construct, which contains only one foldable sGFP domain (Figure 2E, F). Consistent with this interpretation, the reciprocal mCherry-sGFP fusion, in which sGFP occupies the C-terminal position, exhibited no fluorescence enhancement in the presence of hHsp60/10 (Figure 2E, F). Together, these observations suggested that hHsp60/10 may preferentially assist folding of one domain even within the tandem construct.

To directly determine which domain is assisted by hHsp60/10 within the tandem substrate, we introduced the chromophore-inactivating Y66G mutation in either of the two sGFP domains, generating a non-fluorescent variant (NFsGFP) that retains a largely intact β-barrel and therefore reports domain position independently of its own fluorescence (33). Two asymmetric constructs were generated, NFsGFP-sGFP, in which the non-fluorescent domain occupies the N-terminal position, and sGFP-NFsGFP, in which it occupies the C-terminus (Figure 2G). Because only one domain in each construct retains the ability to generate fluorescence, the resulting signal directly identifies which position undergoes productive folding. If hHsp60/10 preferentially assists folding of the N-terminal domain, fluorescence should be observed only when the fluorescent sGFP occupies the N-terminus (Figure 2G).

Consistent with this prediction, upon co-expression with hHsp60/10, fluorescence was enhanced only for sGFP-NFsGFP, in which the fluorescent sGFP domain is located at the N-terminus (Figure 2H). In contrast, placing the non-fluorescent variant at the N-terminus (NFsGFP-sGFP) abolished the fluorescence enhancement with hHsp60/10, despite the presence of an intact, potentially foldable sGFP at the C-terminus (Figure 2H). Moreover, the fluorescence intensity of sGFP-NFsGFP was comparable to that of the original 2x-sGFP construct, confirming that the tandem fluorescence arises entirely from folding of the N-terminal domain (Figure 2H). These results demonstrate that hHsp60/10 preferentially assists folding of the N-terminal domain within a multidomain polypeptide, leaving the C-terminal domain largely unassisted.

To determine whether this positional preference extends beyond the sGFP reporter, we tested MetK, an endogenous GroEL/ES class-III substrate that fails to fold spontaneously (13, 15), fused to yeGFP in either orientation (Figure 2I). Co-expression with hHsp60/10 increased solubility of the fusion protein when MetK occupied the N-terminal position (MetK-yeGFP), but not when it was positioned at the C-terminus (yeGFP-MetK) (Figure 2I, S2B). This observation suggests that the N-terminal positional preference of hHsp60/10 is independent of domain identity and that the human mitochondrial chaperonin engages multidomain substrates in a directional manner.

Because the folding-defective domain was identical across these constructs, and all substrates remained within the size range compatible with GroEL/ES encapsulation, the decline in GroEL/ES assistance with increasing substrate size cannot be attributed to differences in the intrinsic foldability of the reporter or to an inability to encapsulate the substrates. Instead, GroEL/ES assisted the folding as a function of the total size of the polypeptide containing the folding domain, whereas hHsp60/10 assisted the same folding domain irrespective of total size, provided it occupied the N-terminal position. For an identical folding problem, GroEL/ES therefore did not provide domain-autonomous assistance, whereas hHsp60/10 did. This presents hHsp60/10 as the Group I counterpart to the domain-wise (partial) encapsulation of substrates as reported for the Group II chaperonin, TRiC (19), but with a special preference for the N-terminal domain.

### Apical domain tyrosine residues contribute to substrate assistance by hHsp60

We next asked whether this vectorial assistance depends on specific residues of hHsp60. As an initial test of our assay, we examined V72I, a clinically validated loss-of-function variant of hHsp60 that causes hereditary spastic paraplegia 13 (SPG13) (34). Importantly, V72I does not cause any gross biochemical defect in the chaperonin. The mutant complex remains stable and retains ATPase activity, which is mildly elevated relative to wild-type hHsp60 (35, 36). It also shifts the heptamer-tetradecamer equilibrium towards the double-ring (tetradecameric) assembly, which wild-type hHsp60 populates only transiently (36, 37). Despite retaining these biochemical properties, V72I showed reduced folding assistance across all substrates tested in our *in vivo* assay relative to wild-type hHsp60/10 (Figure 3A). Thus, the assay detected the loss of folding activity associated with the clinically established loss-of-function variant of hHsp60, despite the mutant chaperonin remaining stable and catalytically active. This further supports the ability of our *E. coli*-based assay to report genuine hHsp60/10 folding activity, confirming that the reporter detects physiologically relevant chaperonin function.

**Figure 3.**
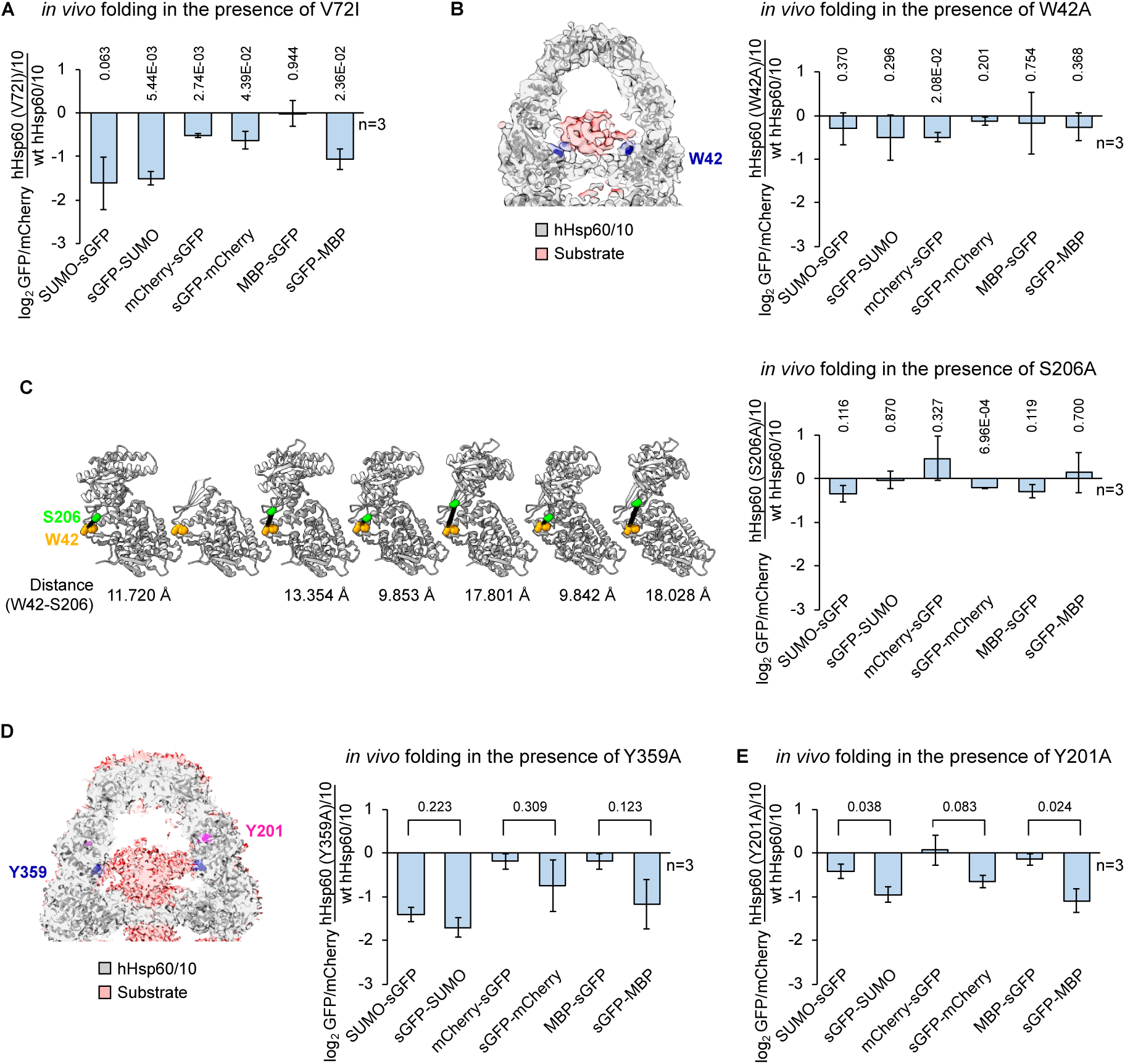
Apical domain residues contribute to substrate assistance by hHsp60/10. **(A**) Effect of the disease-associated V72I mutation on folding of the indicated reporter substrates relative to wild-type hHsp60. **(B)** Left, cryo-EM structure of substrate-bound hHsp60/10 highlighting the interaction between W42 (blue) and the bound substrate (red). The chaperonin is shown in grey. Electron densities were obtained from EMDB:62714 and the corresponding structure from PDB:9L8P (39). Right, effect of W42A substitution on substrate folding relative to wild-type hHsp60/10. **(C)** Left, unwrapped view of the asymmetric hHsp60 heptamer (PDB:8G7M) (37) protomer-specific interactions between W42 (orange) and S206 (green). Cɑ-Cɑ distances between the two residues are indicated for each protomer. Right, effect of S206A substitution on substrate folding relative to wild-type hHsp60/10. **(D)** Left, cryo-EM density (EMDB:29818) and structural model of hHsp60/10 (PDB:8G7O) highlighting residues Y201 (magenta) and Y359 (blue) at the substrate interaction surface (red) (37). Right, effect of Y359A substitutions on substrate folding relative to wild-type hHsp60/10. **(E)** Effect of Y201A substitution on substrate folding relative to wild-type hHsp60/10. Data are mean ± SD from three independent experiments. Statistical significance was determined using two-tailed Student’s t-tests; p-values relative to folding in the presence of wild-type hHsp60/10 are shown above each bar in (A, B and C).

We next attempted to identify which residues contribute to this activity. Recent substrate-bound cryo-EM structures of hHsp60/10 identified W42 in direct contact with the substrate (Figure 3B) (38, 39). Additionally, in the asymmetric ATP-bound heptamer, W42 forms contacts with S206 that differ among protomers (Figure 3C) (37), suggesting that the W42-S206 interaction may itself contribute to the asymmetric distribution of protomer conformations. To understand the contribution of these residues, we created W42A and S206A mutants of hHsp60/10. However, neither mutation altered the assistance across the substrate panel compared with the wild-type chaperonin (Figure 3B, 3C). Therefore, despite the structural proximity of W42 to the substrate and its interaction with S206, disrupting these contacts does not alter folding assistance. The interacting residues identified in these cryo-EM studies were observed in complexes with different substrates, and may therefore be dispensable, or their contributions may be compensated by other interactions during folding of the substrates tested here.

Another cryo-EM structure of hHsp60/10 suggests an interface at the apical domain involving Y201 in the ATP-bound state of hHsp60 and Y359 in the Hsp10/ATP-bound closed conformation as potential substrate-interacting residues (Figure 3D) (37). To understand their contribution to substrate assistance, we generated Y201A and Y359A mutants and tested their activity across our substrate panel. Both mutants showed reduced folding assistance compared with wild-type hHsp60/10 (Figure 3D, E). The reduction was larger for substrates in which sGFP occupied the N-terminal position in the Y201A substitution (Figure 3E) and statistically significant in two of the three constructs and marginally insignificant, with the same trend, in the third construct tested. Y359A substitution also showed a similar trend, but the difference did not reach statistical significance (Figure 3D). These hHsp60 variants express and assemble similarly to the wild-type hHsp60/10, hence the reduced assistance cannot be attributed to a general loss of activity (Figure S3A, B). Additionally, Y201A is not a loss of function mutation like V72I, since it retains its ability to assist SUMO-sGFP, That V72I cannot (Figure 3A, E). Thus, Y201, a substrate acceptor residue at the apical domain of hHsp60, may be important for the positional preference of hHsp60/10 towards N-terminal domains, the details of which remain to be elucidated.

### Translation-coupled folding reveals the N-terminal preference of hHsp60/10

To understand the mechanistic basis of the N-terminal folding preference observed *in vivo*, we first examined whether the distinct substrate preferences of GroEL/ES and hHsp60/10 could be recapitulated *in vitro* using purified components. Chemically denatured sGFP, SUMO-sGFP and MBP-sGFP were allowed to refold in the presence of purified chaperonins and ATP. Consistent with previous reports and our *in vivo* observations, GroEL/ES accelerated refolding of isolated sGFP by ∼5-fold (Figure 4A) (30). This assistance decreased with increasing substrate size, with only a ∼2-fold acceleration observed for SUMO-sGFP refolding and almost no enhancement for MBP-sGFP (Figure 4A). This faithfully recapitulates the substrate size-dependence of GroEL/ES activity observed *in vivo*.

**Figure 4.**
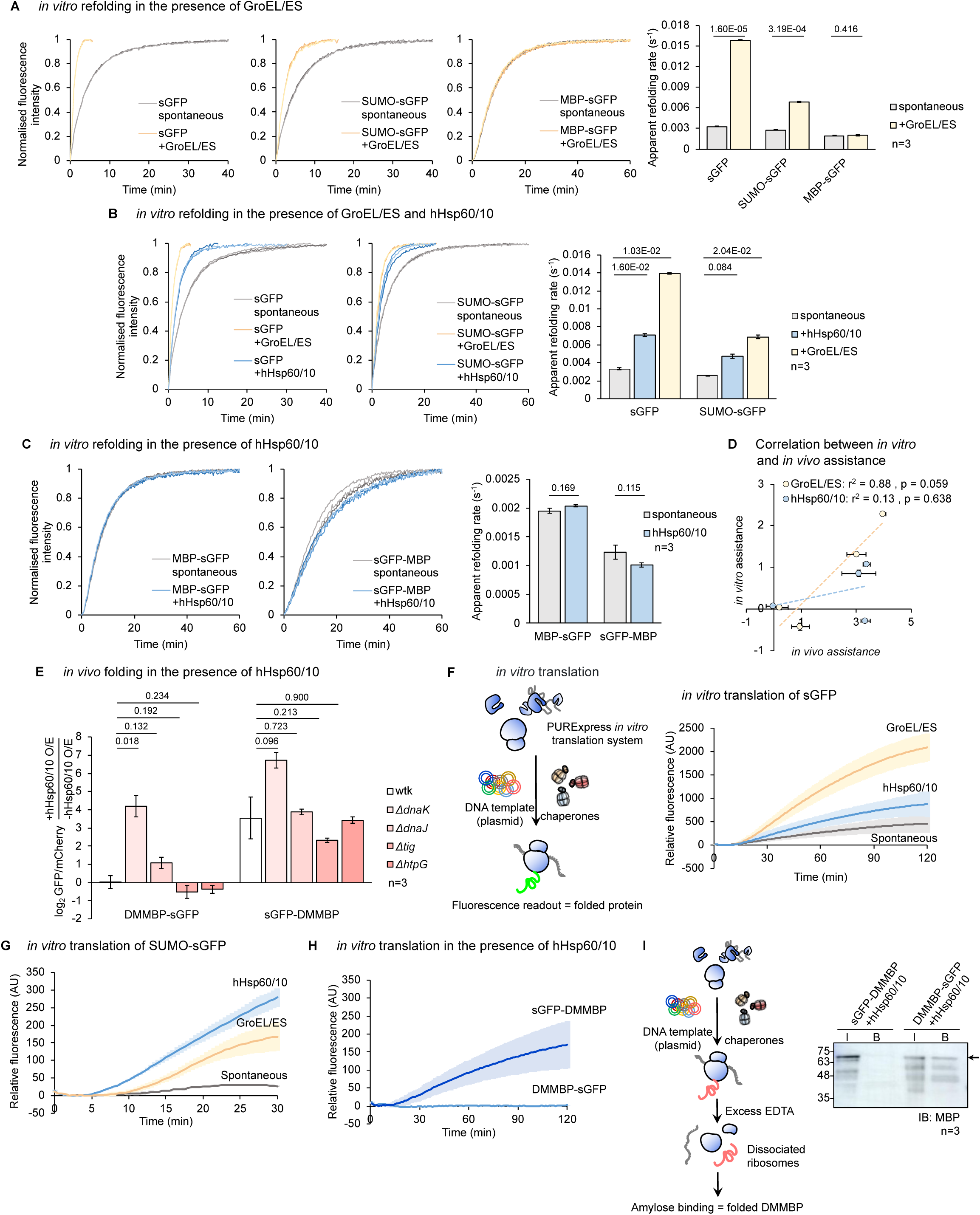
hHsp60/10 promotes vectorial folding of emerging polypeptide chains. **(A)** *In vitro* refolding kinetics of sGFP, SUMO-sGFP and MBP-sGFP in the absence (spontaneous) or presence of GroEL/ES. Right, apparent refolding rates derived from kinetic fits. **(B)** *in vitro* refolding kinetics of sGFP and SUMO-sGFP in the absence (spontaneous) or presence of GroEL/ES or hHsp60/10. Right, apparent refolding rates derived from kinetic fits. **(C)** *in vitro* refolding of MBP-sGFP and sGFP-MBP in the absence (spontaneous) or presence of hhsp60/10. Right, apparent rates of refolding derived from kinetic fits. **(D)** Correlation between *in vitro* and *in vivo* folding assistance of substrates by GroEL/ES or hHsp60/10. Folding assistance is denoted as the log_2 f_old change in the fluorescence (*in vivo*) or apparent refolding rate (*in vitro*) in the presence of chaperonin relative to spontaneous folding. **(E)** Folding of indicated fusion proteins by hHsp60/10 in wild-type (Wtk) and chaperone deletion strains relative to spontaneous folding. **(F)** Left, schematic of translation-coupled folding assay. Purified chaperonin complexes were added to the reconstituted PURExpress transcription-translation system, and GFP fluorescence was monitored in real-time during protein synthesis. Right, translation-coupled folding of sGFP in the absence or presence of GroEL/ES or hHsp60/10. **(G)** Translation coupled folding of SUMO-sGFP measured as in (E). **(H)** Translation-coupled folding of DMMBP-sGFP and sGFP-DMMBP in the presence of hHsp60/10 demonstrating preferential folding of the N-terminal sGFP. **(I)** Left, schematic of assay used to measure DMMBP folding following *in vitro* translation. Translation reactions were terminated with EDTA, folded DMMBP was isolated by amylose affinity chromatography, and recovered protein was analysed by immunoblotting. Right, representative western blot of input (I) and amylose-bound (B) fractions following *in vitro* translation of sGFP-DMMBP and DMMBP-sGFP in the presence of hHsp60/10. Blots were probed with an anti-MBP antibody. Data are mean ± SD from three independent experiments. Statistical significance was determined using two-tailed Student’s t-tests; p-values depicted on top.

By contrast, hHsp60/10 accelerated refolding of sGFP by only ∼2-fold (Figure 4B). Importantly, incorporation of sGFP into the larger SUMO-sGFP construct did not improve hHsp60/10 mediated refolding, with both substrates exhibiting a similar ∼2-fold enhancement (Figure 4B). Likewise, although hHsp60/10 efficiently assisted folding of sGFP-MBP *in vivo*, it failed to enhance refolding of either sGFP-MBP or MBP-sGFP *in vitro* (Figure 4C). This limited activity could not be attributed to defective chaperonin preparations, as purified hHsp60/10 retained its oligomeric assembly and ATPase activity (Figure S4A, B) and its ability to accelerate sGFP refolding marginally. Thus, unlike GroEL/ES, whose *in vivo* folding assistance correlated strongly with its assistance to refolding *in vitro* (Figure 4D), hHsp60/10 assistance to *in vivo* folding did not correlate with its assistance to refolding *in vitro* (Figure 4D). Additionally, *in vitro* refolding reactions with hHsp60/10 failed to recapitulate the N-terminal preference observed in *E.coli* cells (Figure 4C).

To determine whether this discrepancy arose from the absence of cellular factors in the purified system, we repeated the refolding assays using clarified lysates from Wtk cells over-expressing

GroEL/ES or hHsp60/10. Lysates enriched in GroEL/ES enhanced refolding of isolated sGFP but not SUMO-sGFP, whereas lysates enriched in hHsp60/10 failed to enhance refolding of either substrates, recapitulating our observations with the purified chaperonins (Figure S4C). This indicates that the inability to reproduce the *in vivo* assistance of hHsp60/10, *in vitro,* is unlikely to arise solely from compromised functionality of purified chaperonin or absence of cytosolic factors.

The discrepancy between *in vivo* folding and *in vitro* refolding could reflect either a requirement of additional cellular machinery, or a dependence on the manner in which substrates are presented to the chaperonin. We first tested whether hHsp60/10 relies on substrate transfer from other chaperone systems, a mechanism well established in cellular protein biogenesis (40–44). Deletion of the major *E. coli* chaperone systems, DnaK, DnaJ, trigger factor (Tig) or HtpG did not impair hHsp60/10-assisted folding of sGFP-DMMBP, over its assistance to the reciprocal construct DMMBP-sGFP (Figure 4E). Thus, the observed orientation preference for folding assistance does not require an obligatory substrate hand-off from any of these individual cytosolic chaperones.

A key distinction between *in vivo* and *in vitro* experiments is the state of the substrate when it encounters the chaperonin. While denaturation-refolding assays present fully synthesised polypeptides, proteins in cells become available for folding progressively as they emerge from the ribosome. Given the N- to C-terminal direction of protein synthesis and the pronounced preference of hHsp60/10 for N-terminal domains, we hypothesised that may hHsp60/10 preferentially assist folding during protein biogenesis, where N-terminal regions of a polypeptide become accessible before its C-terminal regions. Such translation-associated folding has been recognised as a distinguishing feature of the eukaryotic group-II chaperonin TRiC (20, 45–47) and, more recently, for GroEL/ES (18).

To test whether coupling folding to protein biogenesis could reproduce hHsp60/10 activity, we used the PURE (Protein synthesis Using Recombinant Elements) system, a reconstituted translation system devoid of endogenous *E. coli* chaperones (48). This allowed us to examine the effects of purified GroEL/ES or hHsp60/10 on folding during protein synthesis without contributions from other cellular chaperone systems. Translation reactions were supplemented with purified chaperonins, and GFP fluorescence was monitored as a readout of productive folding (Figure 4F).

Both GroEL/ES and hHsp60/10 enhanced folding of isolated sGFP in the PURE system (Figure 4F). Consistent with both *in vivo* and *in vitro* observations, folding assistance provided by hHsp60/10 remained lower than GroEL/ES, although this observation alone does not distinguish between assistance occurring during translation or post-translationally. Strikingly, however, hHsp60/10 efficiently promoted folding of SUMO-sGFP in the translation reaction, and provided greater assistance than GroEL/ES, reproducing the relative activities observed *in vivo* (Figure 4G). This contrasted with the post-translational refolding experiments in which hHsp60/10 provided only limited assistance to SUMO-sGFP. Thus, coupling folding to protein synthesis was sufficient to recapitulate the substrate preferences of hHsp60/10 observed *in vivo*.

We next asked whether this translation-coupled folding also reproduced the positional preference of hHsp60/10 observed in cells. To this end, we tested reciprocal fusions of sGFP and DMMBP (DMMBP was used instead of MBP for reasons explained in the next paragraph. The assistance to sGFP does not differ between its fusion with MBP or DMMBP *in vivo* (Figure S4D)). Consistent with the N-terminal preference observed *in vivo* (Figure 2A), hHsp60/10 efficiently enhanced folding of sGFP-DMMBP, where sGFP occupies the N-terminal position, during translation-coupled folding (Figure 4H). In contrast, DMMBP-sGFP, where sGFP is at the C-terminus, showed no increase in fluorescence in the presence of hHsp60/10 (Figure 4H). Thus, the N-terminal preference of hHsp60/10 was maintained in the reconstituted translation system.

To determine whether this vectorial folding behaviour extended beyond sGFP, we next examined folding of the DMMBP domain itself. DMMBP is a compromised folder that can fold efficiently with the assistance of GroEL/ES (49). It is a destabilised <u>D</u>ouble <u>M</u>utant of <u>M</u>altose-<u>B</u>inding <u>P</u>rotein and binds amylose only in its native conformation (Figure 4I) (50). Following translation in the presence of purified hHsp60/10, ribosomes were dissociated and the chaperonin activity terminated by addition of excess EDTA (51–53). The reaction mixtures were then incubated with amylose resin, and bound proteins were analysed by immunoblotting to assess DMMBP folding (Figure 4I). DMMBP bound efficiently to amylose when present at the N-terminus of the fusion construct (DMMBP-sGFP), indicating that the domain was folded. In contrast, no amylose binding was detected for sGFP-DMMBP, where DMMBP is at the C-terminus (Figure 4I, S4E). Together with the reciprocal GFP fluorescence measurements, these results demonstrate that the positional preference of hHsp60/10 is not restricted to sGFP. Rather, both folding reporters showed enhanced acquisition of their native conformations when positioned at the N-terminus of a multidomain substrate (Figure 4H, I).

Collectively, these findings suggest that hHsp60/10 mediated folding of multidomain substrates depends on the context in which the polypeptide is presented. Unlike GroEL/ES, hHsp60/10 does not recapitulate it’s *in vivo* substrate preferences during refolding of fully synthesised polypeptides. Instead, its enhanced activity towards multidomain substrates and its preference for N-terminal domains are recapitulated in a reconstituted translation system. These observations support a model in which hHsp60/10 preferentially assists folding of polypeptides as they emerge sequentially during synthesis, rather than acting exclusively on completely synthesised, unfolded proteins. Our assays do not, however, directly demonstrate engagement of hHsp60/10 with a ribosome-attached nascent chain; here, translation-coupled assistance refers to the folding activity observed in the presence of ongoing protein synthesis. Further, because an N-terminal domain is synthesised first and is consequently available for a longer period before synthesis of the C-terminal domain is complete, the observed positional preference could reflect either preferential assistance to N-terminal domains due to their vectorial property (for e.g., peptide dipole) or the kinetics of substrate emergence, the latter being equally consistent with a chaperonin that engages whichever segment becomes accessible first. Distinguishing between these possibilities will require experiments that independently vary the timing of domain emergence independent of sequence orientation.

### hHsp60/10-dependent folding in the mitochondrial matrix retains the N-terminal preference

Although the experiments in *E. coli* and the reconstituted translation system revealed a pronounced preference of hHsp60/10 for N-terminal domains, the physiological relevance of this behaviour remained unclear. In human cells, hHsp60/10 functions within the mitochondrial matrix, where proteins become available for folding through two distinct routes: synthesis by mitochondrial ribosomes and import of nuclear-encoded precursor proteins. To monitor hHsp60/10-dependent folding in the mitochondrial matrix, we sought an *in vivo* folding reporter of chaperonin activity that would be insulated from the defects in mitochondrial translation and protein import that accompany hHsp60 depletion (1, 54–56). We developed a dual-colour folding reporter to distinguish changes in GFP folding from changes in protein expression or mitochondrial targeting. GFP and mCherry were expressed as separate polypeptides from a single mRNA, each carrying an independent, identical mitochondrial matrix-targeting sequence (Figure 5A). The two proteins were therefore targeted independently to the mitochondria, while their overlapping fluorescence marked the mitochondrial network in HeLa cells (Figure 5A). In this system, mCherry served as an internal control for expression and import, allowing GFP folding to be normalised against variations in these processes (Figure 5A).

**Figure 5.**
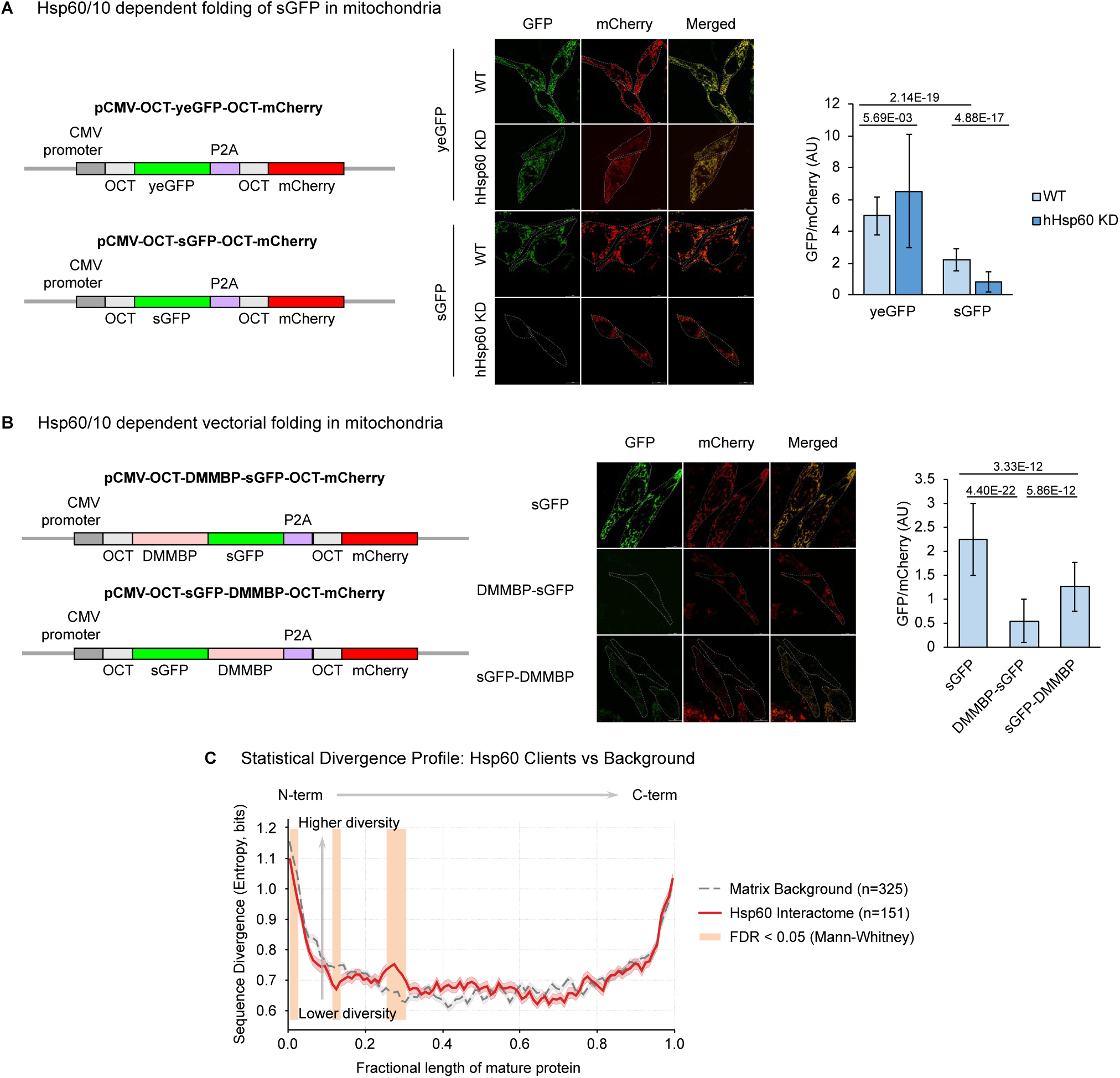
The N-terminal folding preference of hHsp60/10 is preserved in the mitochondrial matrix. **(A)** Schematic of the mitochondrial folding reporter. OCT-targeted GFP variants were co-expressed with OCT-mCherry separated by a P2A peptide to normalise for expression and mitochondrial import. Representative confocal images of HeLa cells expressing mitochondrially targeted yeGFP or sGFP following control or HSPD1 (hHsp60) knock down. Right, quantification of mitochondrial GFP/mCherry fluorescence ratios; n=50. **(B)** Schematic of mitochondrially targeted DMMBP-sGFP and sGFP-DMMBP used to examine the effect of domain order on folding within the mitochondrial matrix. Representative confocal images of HeLa cells expressing the indicated mitochondrial reporter constructs. Right, quantification of mitochondrial GFP/mCherry fluorescence ratios demonstrating preferential folding of substrates carrying sGFP at the N-terminus; n=60. **(C)** Average sequence divergence across metazoans, measured as Shannon entropy, plotted along the normalised length of non-substrate matrix proteins (grey) and the hHsp60 interactome (red). Mitochondrial targeting sequences were excluded before analysis. Shaded regions indicate sequence segments showing significant differences in sequence divergence between the two protein sets. Data are mean ± SD. Statistical significance was determined using two-tailed Student’s t-tests; p-values depicted on top.

Knockdown of hHsp60 did not reduce fluorescence of spontaneously folding yeGFP with respect to the mCherry control (Figure 5A, S5A). In contrast, fluorescence of slow-folding sGFP, which was already substantially lower than that of yeGFP in wild-type cells, decreased further upon hHsp60 depletion (Figure 5B). Importantly, this reduction remained significant after normalisation to mCherry fluorescence, indicating that folding of sGFP within the mitochondrial matrix is reduced on hHsp60 depletion. This assistance recapitulated our observations in *E. coli* cells and in *in vitro* translation coupled folding assays. Matrix-targeted sGFP therefore provides a specific *in vivo* readout of hHsp60/10-mediated folding in mammalian cells.

Having established this reporter, we next asked whether hHsp60 retains its preference for N-terminal domains within the mitochondrial matrix. Reciprocal fusions of sGFP and DMMBP were targeted to mitochondria using the same expression system (Figure 5C). Consistent with our observations in *E. coli*, fluorescence was significantly higher when sGFP occupied the N-terminal position (sGFP-DMMBP) than when it was positioned at the C-terminus (DMMBP-sGFP) (Figure 5B). Furthermore, knockdown of hHsp60 reduced fluorescence from sGFP-DMMBP (Figure S5B), confirming that productive folding of the N-terminal sGFP domain in the matrix depended on hHsp60. Because sGFP-DMMBP and DMMBP-sGFP are identical in mass, this positional difference cannot be attributed to substrate size. Thus, these experiments establish that the N-terminal preference observed in our cellular folding assays is also preserved in the mitochondrial matrix. However, these data do not demonstrate that folding is coupled directly to translocation.

The physiological context of mitochondrial protein biogenesis provides a possible explanation for this preference. Most mitochondrial matrix proteins are encoded in the nucleus, translated in the cytosol and imported through the mitochondrial translocation machinery. Because matrix targeting signals lie predominantly at the N-terminus, precursor proteins enter the matrix vectorially in an N- to C-terminal direction (22–24). Consequently, N-terminal regions become available within the matrix before the remainder of the polypeptide has completed import. The directional preference of hHsp60/10 for N-terminal domains therefore parallels the vectorial presentation of the substrates during mitochondrial import. Such a mechanism could allow the mitochondrial chaperonin to support folding of multidomain proteins without requiring complete encapsulation of the entire polypeptide, as it emerges into the matrix, whether during or shortly after import.

To explore whether physiological hHsp60/10 interactors are consistent with this possibility, we analysed the publicly available hHsp60 interactome dataset (28). Notably, all but one of the reported hHsp60/10 interactors were nuclear-encoded proteins, suggesting that imported precursors constitute a major class of physiological clients for the mitochondrial chaperonin. The same dataset also identified components of the mitochondrial import machinery, including Tim44 and the matrix Hsp70 mortalin (HSPA9), among proteins found in proximity to hHsp60/10. Although these associations are correlative and do not establish direct substrate transfer or folding during import, they raise the possibility that hHsp60/10 engages precursor proteins as they emerge into the matrix during import.

Directly testing the folding of individual domains of endogenous mitochondrial substrates remains challenging because their native conformations cannot be readily monitored *in vivo*. To collect supportive evidence, we therefore asked if the sequences of physiological hHsp60/10 interactors exhibit an evolutionary signature consistent with preferential assistance of N-terminal regions. Molecular chaperones can buffer the destabilising effects of mutations, potentially allowing their substrates to accumulate sequence variations that would otherwise compromise folding (57–63). If hHsp60/10 preferentially assists N-terminal domains, these regions might tolerate more sequence divergence than corresponding regions of mitochondrial matrix proteins that are not identified as hHsp60/10 interactors. Their C-terminal regions, by contrast, should diverge no differently from those non-substrates.

We therefore compared sequence divergence across metazoan orthologues of reported hHsp60/10 substrates and of non-interacting mitochondrial matrix proteins. Because most nuclear-encoded mitochondrial proteins contain an N-terminal mitochondrial targeting sequence (MTS) that is removed upon import, we excluded experimentally annotated or TargetP-predicted targeting sequences before analysing the mature proteins. We additionally excluded the first 20% of each mature sequence to minimise contributions from residual targeting sequences and the unstructured N- and C-terminal regions which do not form part of the final folded proteins. Sequence divergence was then compared across corresponding positions of the mature proteins.

Between the 25th and 40th percentile of the sequence, hHsp60/10 substrates diverged significantly more than non-substrates, whereas their C-terminal halves showed no such difference (Figure 5C, Table S2). This percentile window was identified exploratorily and was not pre-specified; all bins are shown (corrected for multiple comparisons), and the comparison is a difference-of-differences (N-versus C-terminal divergence, interactors versus non-interactors) that cancels per-protein rate determinants such as abundance, although percentile-based mapping is not domain-aware. This enrichment in N-terminal sequence divergence was not explained by increased intrinsic disorder, as the corresponding regions of hHsp60/10 interactors were no more disordered than those of non-interacting matrix proteins (Figure S5C, Table S2). Nor did it reflect differences in the number of residues or number of proteins in each bin between the N- and C-termini (Figure S5D, Table S2).

These observations indicate that N-terminal regions of hHsp60/10 interactors accumulate more mutations over background than their C-terminal halves do. While this does not establish causality and cannot dissociate folding on an N-terminal domain from the C-terminal one, it is consistent with preferential buffering of N-terminal regions by hHsp60/10, relaxing the evolutionary constraints on those regions of its clients.

Together, these findings extend the N-terminal folding preference of hHsp60/10 to its native mitochondrial environment and identify a corresponding pattern of increased sequence divergence among physiological hHsp60 interactors. Along with the translation-coupled folding experiments, they suggest that hHsp60/10 may be adapted to assist multidomain proteins whose domains become accessible sequentially during protein biogenesis. Such a mechanism could enable the mitochondrial chaperonin to support folding of proteins that exceed the geometric constraints of canonical substrate encapsulation.

### Metazoan Hsp60/10 has evolved to assist vectorial folding of larger substrates

To determine when the altered substrate-recognition properties of hHsp60/10 emerged during evolution, we examined homologues of GroEL/ES and Hsp60/10 from representative branches of the tree of life, including prokaryotes (*Cellulomonas flavigena, Candidatus Sulcia muelleri, Sphingomonas wittichii, Acidovorax avenae, Alkalilimnicola ehrlichii, Halorhodospira halophila, Escherichia coli* and *Wigglesworthia brevipalpis*), fungi (*Saccharomyces cerevisiae*), plants (*Cyanidioschyzon merolae* and *Arabidopsis thaliana*), protists (*Giardia intestinalis, Trypanosoma brucei, Tetrahymena thermophila* and *Dictyostelium discoideum*) and metazoans (*Amphimedon queenslandica, Nematostella vectensis, Drosophila melanogaster, Crassostrea gigas, Strongylocentrotus purpuratus* and *Homo sapiens*) (Figure 6A). We first asked whether the size dependence of substrate assistance observed for *E. coli* GroEL/ES was conserved among other prokaryotic homologues. We expressed these chaperonin homologues in Wtk cells alongside our panel of model substrates (Figure S6A). Consistent with our previous observations, the different prokaryotic homologues displayed distinct efficiencies in assisting sGFP folding (Figure 6B, S6B) (30). However, when their activity was compared across substrates of increasing molecular size (sGFP, sGFP-SUMO, sGFP-mCherry and sGFP-MBP), all homologues exhibited a progressive decline in folding assistance with increasing substrate size (Figure 6B). Thus, despite differences in their overall folding efficiencies, the prokaryotic chaperonin homologues retained the size-dependent substrate-assistance pattern characteristic of *E. coli* GroEL/ES.

**Figure 6.**
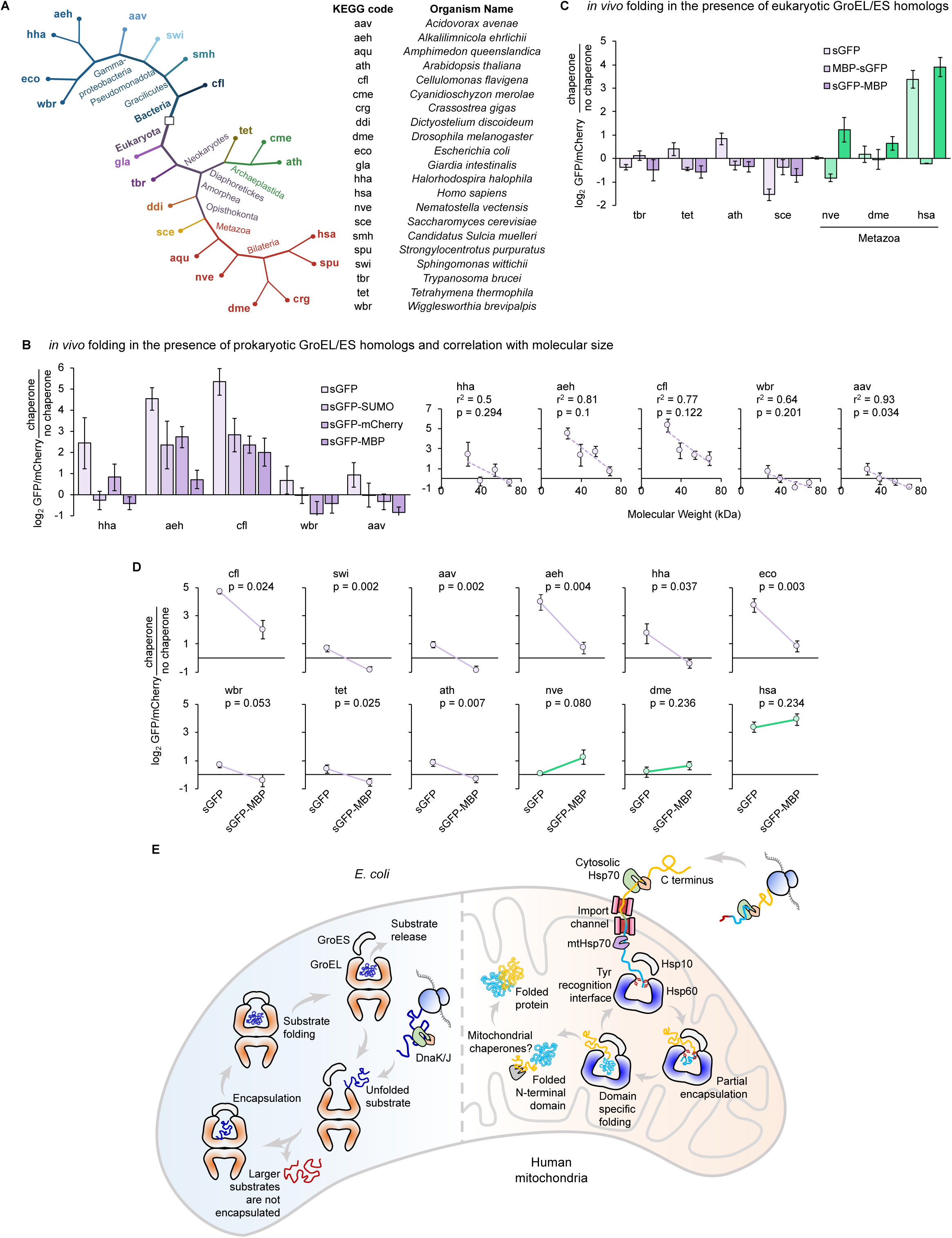
Distinct substrate-assistance properties emerged along the metazoan Hsp60 lineage. **(A)** Representative phylogenetic tree showing the organisms selected from different evolutionary branches to analyse chaperone function. Right, list of the selected organisms and their corresponding 3-letter KEGG codes. **(B)** Left, *in vivo* folding activity of representative prokaryotic GroEL homologs. Right, correlation between chaperone-assisted folding and substrate molecular size. **(C)** *in vivo* folding activity of representative eukaryotic Hsp60/10 homologs. **(D)** Paired dot plots showing the change in folding assistance between sGFP and sGFP-MBP across the indicated chaperone homologs. **(E)** Model summarising the proposed mechanism of substrate engagement and folding by hHsp60 (right), in the context of mitochondrial import-associated folding, alongside the proposed GroEL/ES mechanism (left). Data are mean ± SD from the three independent experiments. Statistical significance was determined using two-tailed Student’s t-tests; p-values depicted on top; correlations were assessed by linear regression.

We next asked whether the N-terminal folding preference of hHsp60/10 was conserved across eukaryotic Hsp60/10 homologues. These homologues were expressed in Wtk cells and tested for their ability to assist folding of isolated sGFP and reciprocal MBP-sGFP and sGFP-MBP fusion proteins. Although we could not establish detectable expression of all the tested homologues in Wtk cells (Figure S6A), those that were successfully expressed showed varying abilities to assist sGFP folding (Figure 6C, S6C). Notably, most of these homologues showed little or no assistance towards either multidomain substrate. However, a subset could efficiently assist sGFP-MBP, in which sGFP occupied the N-terminal position, while showing limited activity towards the reciprocal MBP-sGFP construct (Figure 6C). This positional preference was evident only among the metazoan homologues, resembling the behaviour of human Hsp60/10.

To further distinguish the emergence of positional preference from the broader ability to accommodate larger substrates, we compared the relative folding assistance provided by each homologue towards isolated sGFP and sGFP-MBP. This analysis was restricted to homologues that were detectably expressed and capable of assisting the folding of at least one of the substrates tested, thereby allowing us to compare their ability to accommodate an increase in substrate size. All homologues from prokaryotes, protists, fungi and plants showed a marked reduction in folding assistance towards the larger sGFP-MBP substrate relative to isolated sGFP (Figure 6D). In contrast, metazoan homologues largely retained their ability to assist folding of the larger fusion protein (Figure 6D).

Together, these findings suggest an evolutionary divergence in the substrate-assistance properties of the Hsp60/10 family. Across the prokaryotic, protist, fungal and plant homologues examined, folding assistance remained strongly dependent on substrate size, resembling the behaviour of bacterial GroEL/ES. In contrast, metazoan Hsp60/10 homologues showed an altered substrate preference, with an increased ability to assist larger multidomain substrates and a preference for N-terminal domains. These observations suggest that the ability to accommodate larger substrates and preferentially assist N-terminal domains emerged along the metazoan lineage, indicating a functional divergence of Hsp60/10 from the ancestral size-dependent mode of substrate assistance.

## DISCUSSION

Despite decades of work on how GroEL recognises its clients (64, 65), all current models of group-I chaperonin function are centered on the canonical *E. coli* encapsulation mechanism. Our findings reveal a functional divergence from this mode of substrate assistance in metazoan Hsp60/10. Unlike GroEL/ES, which showed progressively reduced folding assistance with increasing substrate size, human mitochondrial Hsp60/10 efficiently assisted folding within multidomain proteins without a corresponding dependence on their total molecular mass (Figure 2B, C). Importantly, this expanded substrate range could not be explained by an enlarged folding chamber, as the ATP-bound cavity of hHsp60/10 is comparable in volume to that of GroEL/ES (Figure 1I).

The question then shifts from whether a large protein fits inside the cavity to how a vectorially engaged substrate is processed once it is recognised. We hypothesise that the two group-I chaperonins solve the same folding problem through different encapsulation strategies (Figure 6E). GroEL/ES completely encloses a non-native polypeptide within its folding chamber, thereby inherently limiting substrate size by cavity dimensions. Consistent with this, for an identical reporter domain GroEL/ES assistance declined as the flanking chain increased in size, whereas hHsp60/10 assistance did not (Figure 2). This comparison using the same substrate suggests that Hsp60/10 assists in a domain-wise mode, which is analogous to, but architecturally distinct from the domain-wise accommodation described for the Group II chaperonin TRiC (19). One possibility is that hHsp60/10 performs segmental encapsulation of multi-domain substrates: the accessible N-terminal domain is preferentially engaged and accommodated within the folding chamber, while the downstream, still-emerging or C-terminal region remains outside (Figure 6E). However, our data do not distinguish this model from two alternatives that could produce the same positional outcome: a relatively slow-acting chaperonin that preferentially engages whichever segment emerges first (emergence-order kinetics), or folding after import of a precursor held in a recognition-competent state by mitochondrial Hsp70 (hold-then-fold). Nor do our experiments distinguish between cis and trans engagement. Thus, the N-terminal preference and the failure of post-translational refolding are consistent with all these possibilities. Figure 6E presents segmental encapsulation as a mechanistic hypothesis rather than an established mode of substrate engagement. If this model holds true, a chaperonin with a fixed cavity volume could assist the folding of proteins far larger than the ∼60 kDa limit, and the strict N-terminal preference follows naturally. Whether the rest of the chain then folds through successive rounds of engagement with the downstream matrix chaperones, or has evolved to fold spontaneously, remains to be determined.

The functional divergence of hHsp60/10 is consistent with several previously reported differences between the mitochondrial Hsp60 and bacterial GroEL. Functionally, hHsp60 is less effective than GroEL at removing flexibility-driven kinetic traps and at buffering destabilising mutations in model single-domain substrates (30, 66). Structurally, it favours a single-ring complex (67–69), adopts asymmetric ATP-bound states (37, 70), carries metazoan-specific substitutions in its substrate-binding surface (37), and possesses different electrostatic properties at its internal cavity lining (30). The disease-associated V72I mutation provides another indication that hHsp60 function cannot be understood solely through the canonical encapsulation model. Despite retaining complex stability and exhibiting increased oligomeric stability and mildly increased ATP hydrolysis rates (34–36), the mutant bypasses the allosteric changes in conformation as seen in MD simulations (71) and compromises folding assistance (Figure 3A), consistent with a defect in productive engagement; we use V72I as a loss-of-function positive control rather than as evidence against the encapsulation model.

A clue to that mechanism lies in how the chaperonin first encounters its substrate. Most mitochondrial matrix proteins are nuclear-encoded, synthesised on cytosolic ribosomes and imported through the TIM23 translocon, threading into the mitochondrial matrix N-terminus first (22–24, 72, 73). A chaperonin acting during this directional delivery, as proposed for the group-II chaperonin TRiC/CCT (20, 74), would no longer be constrained to encapsulate the whole chain at once, potentially overcoming the size limit imposed by the folding chamber (19, 75). Consistent with this idea, early work placed Hsp60 downstream of mitochondrial Hsp70 during import (1, 42, 76), detected interactions between Hsp60 and translocating chains (77), and reported precursors accumulating upon loss of Hsp60 function (1, 78). However, the field gradually settled on the GroEL paradigm of post-translational refolding (79–81), leaving vectorial substrate engagement largely untested.

Our cellular data are consistent with this picture. Matrix-targeted folding of a stringent reporter depends on hHsp60, and the preference for N-terminal domains is preserved within the matrix (Figure 5B). Further, although correlative, the N-terminal regions (tens of residues after the putative signal sequences) of hHsp60 substrates across metazoan orthologues are more divergent than those of non-substrate matrix proteins, whereas their C-terminal regions are not (Figure 5C), as might be expected if preferential folding of N-terminal regions relaxed the evolutionary constraints upon them. We note, as a correlative and exploratory hypothesis, that the directional preference of the metazoan chaperonin may have consequences for the evolution of its physiological substrates.

What emerges is a possible shift in mechanism of engagement to take care of larger more complex proteins. Rather than expanding the dimensions of its folding chamber, metazoan Hsp60 appears to have expanded its folding repertoire by evolving a fundamentally different, position-dependent mode of substrate assistance that has likely been rebuilt around the directional way its clients are synthesised and delivered.

## LIMITATIONS

Emergence-coupled folding is demonstrated in *E. coli* and in a reconstituted, chaperone-free synthesis system, and these converge on the same positional rule. However, direct engagement of hHsp60/10 with nascent chains on the ribosome in situ, and coupling of folding to translocation, remain to be shown. Our mitochondrial experiments establish hHsp60-dependence and a preserved N-terminal preference within the matrix, but do not by themselves demonstrate that folding is co-translocational. We therefore consider it equally compatible with our data that folding proceeds during translocation, or that the matrix Hsp70 and Hsp40 system may hold imported precursors in a recognition-competent state until the chaperonin acts. Distinguishing between these possibilities will require folding assays coupled directly to import kinetics. We also have no direct structural or biophysical characterisation of the substrate-engaged chaperonin; our attempts were limited by the instability of the purified complex, which also precludes the enclosure assays we apply to GroEL/ES. Further studies are required to unravel the mechanistic complexities of this process.

## ACKNOWLEDGEMENTS

We thank the flow cytometry facility and NGS facility at IGIB, HPC facilities at IGIB and IIIT-D for providing computational resources, and confocal microscopy facility at SNIoE. We thank Rajan Singh for his help at the confocal microscopy facility at SNIoE. K.C. acknowledges grant support from DBT-Wellcome Trust India Alliance. K.M. acknowledges the funding support from Anusandhan National Research Foundation (ANRF), Government of India, for the Advanced Research Grant (ANRF/ARG/2025/003237/LS) and SNIoE core funding. Instrument support was obtained from CSIR and DBT-Wellcome Trust India Alliance. K.M. acknowledges SNIoE for infrastructural support, and K.C. acknowledges CSIR and CSIR-IGIB for infrastructural support. K.C. is a DBT-Wellcome Trust India Alliance Senior Fellow. M.K. and S.R. acknowledge UGC for fellowship support. S.B. acknowledges SNioE and M.K.A. acknowledges IIIT-D for doctoral fellowship.

## AUTHOR CONTRIBUTIONS

|  | M.K | S.R | S.B | P.G | H.T | M.K.A | V.B | A.R | D.K | K.M | K.C |
|---|---|---|---|---|---|---|---|---|---|---|---|
| Conceptualization | 1 |  |  |  |  |  |  |  |  | 1 | 1 |
| Investigation | 1 | 1 | 1 | 1 | 1 | 1 | 1 |  |  |  |  |
| Data Curation | 1 | 1 | 1 | 1 | 1 | 1 | 1 |  |  |  | 1 |
| Methodology | 1 | 1 | 1 |  |  |  |  |  |  |  |  |
| Formal Analysis | 1 | 1 | 1 |  |  |  |  |  |  |  | 1 |
| Validation | 1 |  | 1 |  |  |  |  | 1 |  | 1 | 1 |
| Resources | 1 |  |  |  |  |  |  |  |  |  |  |
| Supervision | 1 |  |  |  |  |  |  | 1 | 1 | 1 | 1 |
| Funding Acquisition |  |  |  |  |  |  |  |  |  | 1 | 1 |
| Project Administration |  |  |  |  |  |  |  |  |  |  | 1 |
| Visualization | 1 |  |  |  |  |  |  |  |  | 1 | 1 |
| Writing - original draft | 1 |  |  |  |  |  |  |  |  |  | 1 |
| Writing - review & editing | 1 | 1 |  |  |  |  |  | 1 | 1 | 1 | 1 |

All authors have read the manuscript.

## DECLARATION OF THE USE OF AI

The authors have used generative AI to edit the manuscript and structure it at places. At no point however, the AI has been used to replace critical thinking and framing of the scientific questions.

## REFERENCES

1. M. Y. Cheng et al., Mitochondrial heat-shock protein hsp60 is essential for assembly of proteins imported into yeast mitochondria. Nature 337, 620–625 (1989).

2. O. Fayet, T. Ziegelhoffer, C. Georgopoulos, The groES and groEL heat shock gene products of Escherichia coli are essential for bacterial growth at all temperatures. Journal of bacteriology 171, 1379–1385 (1989).

3. F. U. Hartl, A. Bracher, M. Hayer-Hartl, Molecular chaperones in protein folding and proteostasis. Nature 475, 324–332 (2011).

4. M. Hayer-Hartl, A. Bracher, F. U. Hartl, The GroEL–GroES chaperonin machine: a nano-cage for protein folding. Trends in biochemical sciences 41, 62–76 (2016).

5. A. L. Horwich, W. A. Fenton, E. Chapman, G. W. Farr, Two families of chaperonin: physiology and mechanism. Annu. Rev. Cell Dev. Biol. 23, 115–145 (2007).

6. D. S. Reading, R. L. Hallberg, A. M. Myers, Characterization of the yeast HSP60 gene coding for a mitochondrial assembly factor. Nature 337, 655–659 (1989).

7. M. Mayhew et al., Protein folding in the central cavity of the GroEL–GroES chaperonin complex. Nature 379, 420–426 (1996).

8. A. L. Horwich, G. W. Farr, W. A. Fenton, GroEL− GroES-mediated protein folding. Chemical reviews 106, 1917–1930 (2006).

9. J. S. Weissman et al., Mechanism of GroEL action: productive release of polypeptide from a sequestered position under GroES. Cell 83, 577–587 (1995).

10. Z. Xu, A. L. Horwich, P. B. Sigler, The crystal structure of the asymmetric GroEL–GroES–(ADP) 7 chaperonin complex. Nature 388, 741–750 (1997).

11. B.-W. Ying, H. Taguchi, M. Kondo, T. Ueda, Co-translational involvement of the chaperonin GroEL in the folding of newly translated polypeptides. Journal of Biological Chemistry 280, 12035–12040 (2005).

12. C. Sakikawa, H. Taguchi, Y. Makino, M. Yoshida, On the maximum size of proteins to stay and fold in the cavity of GroEL underneath GroES. Journal of Biological Chemistry 274, 21251–21256 (1999).

13. Y.-C. Tang et al., Structural features of the GroEL-GroES nano-cage required for rapid folding of encapsulated protein. Cell 125, 903–914 (2006).

14. K. Fujiwara, Y. Ishihama, K. Nakahigashi, T. Soga, H. Taguchi, A systematic survey of in vivo obligate chaperonin-dependent substrates. The EMBO journal 29, 1552 (2010).

15. M. J. Kerner et al., Proteome-wide analysis of chaperonin-dependent protein folding in Escherichia coli. Cell 122, 209–220 (2005).

16. T. K. Chaudhuri, G. W. Farr, W. A. Fenton, S. Rospert, A. L. Horwich, GroEL/GroES-mediated folding of a protein too large to be encapsulated. Cell 107, 235–246 (2001).

17. V. Dahiya, T. K. Chaudhuri, Functional intermediate in the refolding pathway of a large and multidomain protein malate synthase G. Biochemistry 52, 4517–4530 (2013).

18. A. Roeselová et al., GroEL/ES chaperonin unfolds then encapsulates a nascent protein on the ribosome. Nature Communications 16, 9976 (2025).

19. F. Rüßmann et al., Folding of large multidomain proteins by partial encapsulation in the chaperonin TRiC/CCT. Proceedings of the National Academy of Sciences 109, 21208–21215 (2012).

20. J. Frydman, E. Nimmesgern, K. Ohtsuka, F. U. Hartl, Folding of nascent polypeptide chains in a high molecular mass assembly with molecular chaperones. Nature 370, 111–117 (1994).

21. W. J. Netzer, F. U. Hartl, Recombination of protein domains facilitated by co-translational folding in eukaryotes. Nature 388, 343–349 (1997).

22. A. Chacinska, C. M. Koehler, D. Milenkovic, T. Lithgow, N. Pfanner, Importing mitochondrial proteins: machineries and mechanisms. Cell 138, 628–644 (2009).

23. W. Neupert, J. M. Herrmann, Translocation of proteins into mitochondria. Annu. Rev. Biochem. 76, 723–749 (2007).

24. M. Schleyer, W. Neupert, Transport of proteins into mitochondria: translocational intermediates spanning contact sites between outer and inner membranes. Cell 43, 339–350 (1985).

25. K. Chakraborty et al., Chaperonin-catalyzed rescue of kinetically trapped states in protein folding. Cell 142, 112–122 (2010).

26. Z. Lin, D. Madan, H. S. Rye, GroEL stimulates protein folding through forced unfolding. Nature structural & molecular biology 15, 303–311 (2008).

27. M. J. Todd, G. H. Lorimer, D. Thirumalai, Chaperonin-facilitated protein folding: optimization of rate and yield by an iterative annealing mechanism. Proceedings of the National Academy of Sciences 93, 4030–4035 (1996).

28. A. S. Bie et al., An inventory of interactors of the human HSP60/HSP10 chaperonin in the mitochondrial matrix space. Cell Stress and Chaperones 25, 407–416 (2020).

29. A. Sadat et al., GROEL/ES buffers entropic traps in folding pathway during evolution of a model substrate. Journal of molecular biology 432, 5649–5664 (2020).

30. A. Sadat et al., Conserved and divergent chaperoning effects of Hsp60/10 chaperonins on protein folding landscapes. Proceedings of the National Academy of Sciences 119, e2118465119 (2022).

31. H. Fan, A. E. Mark, Mimicking the action of GroEL in molecular dynamics simulations: application to the refinement of protein structures. Protein science 15, 441–448 (2006).

32. D.-H. Chen, J.-L. Song, D. T. Chuang, W. Chiu, S. J. Ludtke, An expanded conformation of single-ring GroEL-GroES complex encapsulates an 86 kDa substrate. Structure 14, 1711–1722 (2006).

33. D. P. Barondeau, C. D. Putnam, C. J. Kassmann, J. A. Tainer, E. D. Getzoff, Mechanism and energetics of green fluorescent protein chromophore synthesis revealed by trapped intermediate structures. Proceedings of the National Academy of Sciences 100, 12111–12116 (2003).

34. J. J. Hansen et al., Hereditary spastic paraplegia SPG13 is associated with a mutation in the gene encoding the mitochondrial chaperonin Hsp60. The American Journal of Human Genetics 70, 1328–1332 (2002).

35. P. Bross, R. Magnoni, A. Sigaard Bie, Molecular chaperone disorders: defective Hsp60 in neurodegeneration. Current Topics in Medicinal Chemistry 12, 2491–2503 (2012).

36. L. Chen, A. Syed, A. Balaji, Hereditary spastic paraplegia SPG13 mutation increases structural stability and ATPase activity of human mitochondrial chaperonin. Scientific Reports 12, 18321 (2022).

37. J. R. Braxton, H. Shao, E. Tse, J. E. Gestwicki, D. R. Southworth, Asymmetric apical domain states of mitochondrial Hsp60 coordinate substrate engagement and chaperonin assembly. Nature Structural & Molecular Biology 31, 1848–1858 (2024).

38. K. Ehses et al., Structural remodeling of the mitochondrial protein biogenesis machinery under proteostatic stress. Science Advances 12, eaed3579 (2026).

39. M. Jung, M. Kim, S. J. Ham, J. Chung, S.-H. Roh, In situ characterization of mitochondrial Hsp60-Hsp10 chaperone complex under folding stress. Science advances 11, eadw6064 (2025).

40. D. Balchin, M. Hayer-Hartl, F. U. Hartl, In vivo aspects of protein folding and quality control. Science 353, aac4354 (2016).

41. T. Langer et al., Successive action of DnaK, DnaJ and GroEL along the pathway of chaperone-mediated protein folding. Nature 356, 683–689 (1992).

42. U. Manning-Krieg, P. Scherer, G. Schatz (1991) Sequential actionof mitochondrial chaperones in protein import into the matrix. EMBO (Eur. Mol. Biol. Organ.) J. 10: 3273–3280. (Manning-Krieg327310EMBO (Eur. Mol. Biol. Organ.) J).

43. R. Rosenzweig, N. B. Nillegoda, M. P. Mayer, B. Bukau, The Hsp70 chaperone network. Nature reviews molecular cell biology 20, 665–680 (2019).

44. R. Wadhwa et al., Identification and characterization of molecular interactions between mortalin/mtHsp70 and HSP60. Biochemical Journal 391, 185–190 (2005).

45. J. Frydman, Folding of newly translated proteins in vivo: the role of molecular chaperones. Annual review of biochemistry 70, 603–647 (2001).

46. V. Thulasiraman, C. F. Yang, J. Frydman, In vivo newly translated polypeptides are sequestered in a protected folding environment. The EMBO journal 18, 85–95 (1999).

47. A. Y. Yam et al., Defining the TRiC/CCT interactome links chaperonin function to stabilization of newly made proteins with complex topologies. Nature structural & molecular biology 15, 1255–1262 (2008).

48. Y. Shimizu et al., Cell-free translation reconstituted with purified components. Nature biotechnology 19, 751–755 (2001).

49. J. D. Wang, M. D. Michelitsch, J. S. Weissman, GroEL-GroES-mediated protein folding requires an intact central cavity. Proceedings of the National Academy of Sciences 95, 12163–12168 (1998).

50. S.-Y. Chun, S. Strobel, P. Bassford Jr, L. Randall, Folding of maltose-binding protein. Evidence for the identity of the rate-determining step in vivo and in vitro. Journal of Biological Chemistry 268, 20855–20862 (1993).

51. G. Blobel, Release, identification, and isolation of messenger RNA from mammalian ribosomes. Proceedings of the National Academy of Sciences 68, 832–835 (1971).

52. D. J. Klein, P. B. Moore, T. A. Steitz, The contribution of metal ions to the structural stability of the large ribosomal subunit. Rna 10, 1366–1379 (2004).

53. D. Sabatini, Y. Tashiro, G. Palade, On the attachment of ribosomes to microsomal membranes. Journal of molecular biology 19, 503-IN509 (1966).

54. A. S. Bie et al., A cell model to study different degrees of Hsp60 deficiency in HEK293 cells. Cell Stress and Chaperones 16, 633–640 (2011).

55. J. H. Christensen et al., Inactivation of the hereditary spastic paraplegia-associated Hspd1 gene encoding the Hsp60 chaperone results in early embryonic lethality in mice. Cell Stress and Chaperones 15, 851–863 (2010).

56. C. Cömert et al., HSP60 chaperone deficiency disrupts the mitochondrial matrix proteome and dysregulates cholesterol synthesis. Molecular Metabolism 88, 102009 (2024).

57. D. Bogumil, T. Dagan, Chaperonin-dependent accelerated substitution rates in prokaryotes. Genome biology and evolution 2, 602–608 (2010).

58. S. Maisnier-Patin et al., Genomic buffering mitigates the effects of deleterious mutations in bacteria. Nature genetics 37, 1376–1379 (2005).

59. C. Queitsch, T. A. Sangster, S. Lindquist, Hsp90 as a capacitor of phenotypic variation. Nature 417, 618–624 (2002).

60. S. L. Rutherford, S. Lindquist, Hsp90 as a capacitor for morphological evolution. Nature 396, 336–342 (1998).

61. N. Tokuriki, D. S. Tawfik, Chaperonin overexpression promotes genetic variation and enzyme evolution. Nature 459, 668–673 (2009).

62. T. A. Williams, M. A. Fares, The effect of chaperonin buffering on protein evolution. Genome biology and evolution 2, 609–619 (2010).

63. K. T. Wyganowski, M. Kaltenbach, N. Tokuriki, GroEL/ES buffering and compensatory mutations promote protein evolution by stabilizing folding intermediates. Journal of Molecular Biology 425, 3403–3414 (2013).

64. K. Mapa, S. Tiwari, V. Kumar, G. G. Jayaraj, S. Maiti, Information encoded in non-native states drives substrate-chaperone pairing. Structure 20, 1562–1573 (2012).

65. J. Martin et al., Chaperonin-mediated protein folding at the surface of groEL through a’molten globule’-like intermediate. Nature 352, 36–42 (1991).

66. D. Gautam, M. Aaquib, M. Kochar, K. Chakraborty, Dicarboxylic acids synergize with yeast and human Hsp60/10 systems to mimic GroEL/ES. bioRxiv, 2024.2005. 2015.594092 (2024).

67. D. P. Klebl et al., Cryo-EM structure of human mitochondrial HSPD1. Iscience 24 (2021).

68. K. L. Nielsen, N. J. Cowan, A single ring is sufficient for productive chaperonin-mediated folding in vivo. Molecular cell 2, 93–99 (1998).

69. S. Vilasi et al., Human Hsp60 with its mitochondrial import signal occurs in solution as heptamers and tetradecamers remarkably stable over a wide range of concentrations. PloS one 9, e97657 (2014).

70. S. Nisemblat, O. Yaniv, A. Parnas, F. Frolow, A. Azem, Crystal structure of the human mitochondrial chaperonin symmetrical football complex. Proceedings of the National Academy of Sciences 112, 6044–6049 (2015).

71. L. Torielli et al., Pathogenic mutation impairs functional dynamics of Hsp60 in mono-and oligomeric states. Nature Communications 16, 3158 (2025).

72. W. Voos, Chaperone–protease networks in mitochondrial protein homeostasis. Biochimica et Biophysica Acta (BBA)-Molecular Cell Research 1833, 388–399 (2013).

73. N. Wiedemann, N. Pfanner, Mitochondrial machineries for protein import and assembly. Annual review of biochemistry 86, 685–714 (2017).

74. J. Frydman, F. U. Hartl, Principles of chaperone-assisted protein folding: differences between in vitro and in vivo mechanisms. Science 272, 1497–1502 (1996).

75. J. J. Kelly et al., Snapshots of actin and tubulin folding inside the TRiC chaperonin. Nature structural & molecular biology 29, 420–429 (2022).

76. J. Ostermann, A. Horwich, W. Neupert, F.-U. Hartl, Protein folding in mitochondria requires complex formation with hsp60 and ATP hydrolysis. Nature, 125–130 (1989).

77. H. Koll et al., Antifolding activity of hsp60 couples protein import into the mitochondrial matrix with export to the intermembrane space. Cell 68, 1163–1175 (1992).

78. E. M. Hallberg, Y. Shu, R. L. Hallberg, Loss of mitochondrial hsp60 function: nonequivalent effects on matrix-targeted and intermembrane-targeted proteins. Molecular and Cellular Biology 13, 3050–3057 (1993).

79. A. P. Ben-Zvi, J. Chatellier, A. R. Fersht, P. Goloubinoff, Minimal and optimal mechanisms for GroE-mediated protein folding. Proceedings of the National Academy of Sciences 95, 15275–15280 (1998).

80. K. L. Ewalt, J. P. Hendrick, W. A. Houry, F. U. Hartl, In vivo observation of polypeptide flux through the bacterial chaperonin system. Cell 90, 491–500 (1997).

81. D. J. Hartman, B. P. Surin, N. E. Dixon, N. J. Hoogenraad, P. B. Høj, Substoichiometric amounts of the molecular chaperones GroEL and GroES prevent thermal denaturation and aggregation of mammalian mitochondrial malate dehydrogenase in vitro. Proceedings of the National Academy of Sciences 90, 2276–2280 (1993).

