## Supplemental methods and figures for "First in, first folded: evolution of vectorial substrate folding by metazoan Hsp60/10"

**SUPPLEMENTAL INFORMATION**

**METHODS**

**Strains, plasmids and proteins**

*E. coli* DH5α was used for molecular cloning, Wild-type *E. coli* K-12 strain BW25113 (hereafter referred to as Wtk strain) was used for *in vivo* experiments. Proteins were expressed in either BL21(DE3) Rosetta or WT cells, as indicated for individual experiments. Gene deletion strains were obtained from *E. coli* Genetic Stock Center (CGSC) as part of the Keio collection (82).

**Protein purification**

*Purification of sGFP*

BL21 (DE3) Rosetta cells transformed with pET-Smt3-sGFP encoding $His_{10}$-SUMO-sGFP were grown in LB medium supplemented with 50 µg/mL kanamycin to an $OD_{600}$ of ~0.8. Protein expression was induced with 0.5 mM IPTG and cultures were incubated overnight at 18 °C. Cells were harvested and lysed by sonication in Buffer A (1x PBS, 5 mM imidazole, 5% glycerol) containing 2 mM PMSF and 1x Protease Inhibitor Cocktail (PIC). Lysate was clarified by centrifugation (40,000 x g, 20 min, 4 °C) and the soluble fraction was incubated with Ni-NTA agarose for 3 h at 4 °C. After washing with 10 column volumes (CV) of Buffer A, bound protein was incubated overnight with 10 µg/mL of His-tagged ULP protease in 3 CV of Buffer A to remove the $His_{10}$-SUMO tag. Cleaved sGFP was collected, concentrated and exchanged into a storage buffer (1x PBS, 2 mM DTT and 5% glycerol). Protein concentration estimated using BCA assay kit (Thermo Scientific #23225), and purified protein was stored at -80 °C.

*Purification of SUMO-sGFP*

$His_{10}$-SUMO-sGFP was expressed as described above. Following binding of clarified lysate to Ni-NTA agarose and washing with Buffer A, the protein was eluted with Buffer A supplemented with 300 mM imidazole, dialysed overnight at 4 °C to remove imidazole, concentrated and stored in 1x PBS containing 2 mM DTT at -80 °C.

*Purification of MBP-sGFP and sGFP-MBP*

BL21 (DE3) Rosetta transformed with pBAD plasmids encoding MBP-sGFP or sGFP-MBP were grown in LB media containing 100 µg/mL ampicillin to an $OD_{600}$ of ~0.8. Protein expression was induced with 0.2% (w/v) arabinose and cultures were incubated overnight at 18 °C. Cells were lysed by sonication in

Buffer B (1x PBS, 1 mM EDTA, 1 mM DTT) containing 2 mM PMSF, 1x PIC and 10 µg/mL lysozyme. Clarified lysates were applied to an amylose column. Following washing with 10 CV of Buffer B, bound proteins were eluted with 3 CV of Buffer B containing 10 mM maltose. Eluted proteins were dialysed overnight at 4 °C to remove maltose and EDTA, concentrated, snap-frozen in liq. $N_2$ and stored in 1x PBS containing 2 mM DTT at -80 °C.

*Purification of GroEL and human Hsp60*

GroEL and hHsp60 were purified from Wtk cells transformed with pOFX-GroEL/ES or pOFX-human Hsp60/10, respectively. Cells were grown in LB medium containing 35 µg/mL chloramphenicol to an $OD_{600}$ of 0.8-1.0, induced with 1 mM IPTG for 3 h at 37 °C, and lysed by sonication in Buffer C (25 mM Tris-Cl pH 8.0, 5 mM EDTA, 2 mM DTT) supplemented with 2 mM PMSF and 10 µg/mL lysozyme. Clarified lysates were adjusted to 20% $(NH_4)_2SO_4$ and loaded onto a Butyl Sepharose column pre-equilibrated in Buffer C containing 20% $(NH_4)_2SO_4$. After washing with Buffer C containing 20% $(NH_4)_2SO_4$ and 20% methanol, proteins were eluted using a step gradient from 20% to 0% $(NH_4)_2SO_4$ in Buffer C. Fractions containing GroEL or Hsp60 were concentrated and further purified by size-exclusion chromatography on a Superdex 200 pg 16/60 column equilibrated in Buffer D (25 mM Tris-Cl pH 7.4, 150 mm KCl, 10 mM $MgCl_2$ and 2 mM DTT). Fractions containing GroEL or Hsp60 were concentrated, snap-frozen in liquid $N_2$ and stored at -80 °C.

*Purification of GroES and human Hsp10*

GroES and human Hsp10 were purified from the corresponding *E. coli* expression strains as described above. Following cell lysis and clarification, clarified lysates were applied to a DEAE Sepharose column equilibrated in Buffer C. Proteins were eluted using a step gradient of 100 mM- 1 M NaCl in Buffer C. Fractions containing GroES or hHsp10 were further purified by size-exclusion chromatography on a Superdex 200 pg 16/60 column equilibrated in Buffer D. Fractions containing GroES or Hsp10 were concentrated, snap-frozen in liquid $N_2$ and stored at -80 °C.

**Spontaneous and chaperone assisted *in vitro* refolding assays**

*Purified chaperone based refolding assay*

20 µM of purified sGFP was denatured in 6M guanidine hydrochloride (GuHCl) prepared in Buffer E (25 mM Tris-Cl pH 7.4, 20 mM KCl, 10 mM $MgCl_2$, 2 mM DTT) for 1 h at 25 °C. Refolding was initiated by 100-fold dilution into Buffer E, yielding a final substrate concentration of 200 nM. Chaperone-assisted refolding reactions contained 400 nM tetradecameric GroEL or Hsp60 together with 800 nM heptameric

GroES or Hsp10, corresponding to a substrate:GroEL:GroES molar ratio of 1:2:4, sufficient to ensure complete substrate encapsulation. Reactions were initiated by the addition of 2 mM ATP (30).

Refolding was monitored at 25 °C by measuring GFP fluorescence using a Fluorolog-3 spectrofluorometer (Horiba) with excitation at 488 nm (slit width 2 nm) and emission at 515 nm (slit width 5 nm). Kinetic traces were fitted to a single-exponential function in Origin to determine apparent refolding rate constants.

SUMO-sGFP refolding assays were performed under identical conditions, except that 12.5 μM substrate was denatured prior to dilution, yielding a final substrate concentration of 125 nM. Chaperone concentrations were adjusted proportionally to 250 nM tetradecameric GroEL or Hsp60 and 500 nM heptameric GroES or Hsp10 to maintain the same substrate:GroEL:GroES stoichiometry. Where indicated, 250 nM of ULP protease was included in the refolding reaction.

*Whole-cell lysate based refolding assay*

Wtk cells over-expressing GroEL/ES or hHsp60/10 were grown and induced as described above. Cells were harvested and lysed by sonication in a buffer containing 25 mM Tris-Cl pH 7.4, 200 mM KCl, 5 mM EDTA, 2 mM DTT, 2 mM PMSF and 10 μg/mL lysozyme. Lysates were clarified and diluted 4-fold into Buffer E supplemented with 2 mM ATP. Refolding reactions were initiated by addition of denatured substrates to final concentrations of 200 nM sGFP or 125 nM SUMO-sGFP. GFP fluorescence was monitored as described above, and apparent refolding rates were determined by single-exponential fitting.

**Fluorescence anisotropy**

200 nM of unfolded sGFP was added to Buffer E containing 400 nM of purified GroEL. ATP·$AlF_x$ was generated by adding 2 mM ATP, 2 mM $AlCl_3$ and 12 mM NaF, followed by addition of 800 nM of GroES for assembly of GroEL-substrate-GroES complexes and incubated at 25 °C for 30 min before measurement. Fluorescence anisotropy was recorded using a Fluorolog 3 spectrofluorometer (Horiba) with excitation at 488nm (slit width 2nm) and emission at 515nm (slit width 5nm) at 25 °C. For SUMO-sGFP, 125 nM of denatured protein, 250 nM of GroEL and 500 nM of GroES were used.

**Chaperone assisted *in vivo* folding assays**

Wtk cells were co-transformed with pBAD plasmids encoding GFP substrate constructs and pOFX plasmids encoding GroEL/ES, hHsp60/10 or the indicated chaperonin homologs. Overnight cultures were diluted into LB medium supplemented with 100 μg/mL ampicillin and 35μg/mL chloramphenicol and grown at 37 °C to an $OD_{600}$ of ~0.6. Chaperonin expression was induced with 1 mM IPTG for 30 min, followed by induction of GFP substrate expression with 0.2% (w/v) arabinose at 37 °C for 3 hours. Cells were diluted in PBS and incubated for a further 30 min at 37 °C to allow maturation of mCherry fluorophore before analysis by flow cytometry using a BD FACSymphony A1 cytometer. GFP folding efficiency was quantified as the median cellular GFP/mCherry fluorescence ratio.

**Solubility assay**

BL21 (DE3) Rosetta cells expressing MetK-yeGFP or yeGFP-MetK were lysed in Buffer F (1x PBS, 2 mM DTT, 2 mM PMSF, 1x PIC and 1 mg/mL lysozyme) by ten freeze-thaw cycles. Lysates were clarified by centrifugation (20,000 x g, 20 min, 4 °C). Equivalent amounts of total soluble protein (30 µg) and corresponding insoluble fractions were resolved by SDS-PAGE.

**Translation-coupled *in vitro* folding assays**

Translation reactions were assembled using PURExpress *in vitro* protein synthesis kit (NEB #E6800) according to the manufacturer's protocol with minor modifications. Reactions were supplemented with 100 U/mL SUPERase·In RNase inhibitor, 2 mM DTT, 250 nM GroEL or hHsp60 tetradecamer, 500 nM GroES or hHsp10 heptamer and 2 mM ATP. 50 ng plasmid DNA encoding the indicated substrate was used as the template. Reactions were initiated by incubation at 37 °C.

*GFP-based folding assays*

Folding of GFP was monitored continuously by measuring GFP fluorescence in real time using a BioRad Opus 384 plate reader. Fluorescence traces were analysed using CFX Maestro software.

*DMMBP folding assays*

For DMMBP fused constructs, reactions were incubated for 90 min at 37 °C before termination by addition of EDTA to a final concentration of 20 mM to dissociate ribosomes and prevent further translation. Samples were diluted 10-fold into Buffer F (25 mM Tris-Cl pH 7.4, 20 mM KCl, 20 mM EDTA, 2 mM DTT) and incubated with amylose resin for 30 min at 25 °C to capture folded DMMBP. After washing, bound proteins were eluted with Buffer F containing 10 mM maltose and analysed by western blotting using an anti-MBP antibody.

**Enzyme coupled ATP hydrolysis assay**

ATP hydrolysis was measured using an enzyme coupled ATP consumption assay. Reactions containing 2 mM phosphoenol pyruvate, 20/30 U/mL pyruvate kinase/ lactate dehydrogenase (PK/LDH) enzyme mix,

1 mM NADH and 1 mM ATP in Buffer D were pre-incubated for 5 min at 25 °C to remove contaminating ADP before addition of 2 μM heptameric GroES or hHsp10. ATP hydrolysis was initiated by addition of 1 μM tetradecameric GroEL or hHsp60. NADH oxidation was monitored at 340 nm using a Tecan Infinite M200 Pro plate reader, and ATPase rates were calculated from the rate of NADH consumption and expressed as μM ATP hydrolysed per minute per μM of tetradecameric chaperonin (83, 84).

**Mammalian cell culture and transfection**

HeLa cells were maintained in Dulbecco's modified Eagle medium (DMEM) supplemented with 10% fetal bovine serum (FBS; Gibco) and 1% penicillin-streptomycin at 37 °C in a humidified incubator with 5% $CO_2$. Plasmid DNA encoding mitochondria targeted substrates and HSPD1 targeting siRNA were transfected using Lipofectamine 2000 TM (Invitrogen) according to the manufacturer's protocol. HSPD1 knockdown was verified by western blotting using antibodies against Hsp60, with GAPDH serving as a loading control.

**Live cell imaging and image analysis**

For imaging experiments, cells were seeded onto 35 mm glass-bottom dishes (SPL Life Sciences) and cultured for 24 h before transfection with plasmids and siRNA as described above. Live-cell imaging was performed on a Nikon AXR NSPARC MP + Ti-E confocal microscope equipped with advanced GASP detectors using an Apochromat 100X 1.4 NA oil-immersion objective. Imaging was performed at 37 °C under humidified 5% $CO_2$. GFP and mCherry were excited using 488 nm and 560 nm lasers, respectively. Mean cellular fluorescence intensity was quantified by manually defining regions of interest (ROI) using Nikon NIS-Elements software.

**Quantification and statistical analysis**

Statistical analyses were performed using MATLAB and Origin unless otherwise indicated. Flow cytometry data were analysed using in-house MATLAB scripts. Unless otherwise indicated, statistical comparisons were performed using two-tailed Student's t-tests, and correlations were assessed using Spearman's rank correlation coefficient. Statistical tests, sample sizes, and definitions of error bars are indicated in the corresponding figure legends.

**Computational methods**

***Comparison of size of metazoan mitochondrial proteome with the E. coli proteome***

Metazoan mitochondrial protein IDs and sequences were obtained from MMPdb (https://mmpdb.eeob.iastate.edu/) along with their MTS cleavage sites. The mature sequence was derived using the cleavage site proposed by the database. The mitochondrial protein sequence lengths were then compared between the whole *E. coli* proteome (available through UNIPROT). Mann-Whitney U test (also known as Wilcoxon rank-sum test) along with Benjamini-Hochberg correction for multiple comparison was used to obtain the p-values.

***Comparison of the size limit of GroEL and Hsp60 interactome***

*Molecular weight determination*

Molecular weights of GroEL substrates were taken as reported (mass_kDa, groel_substrates_standardized.tsv). For all other protein sets, molecular weight was calculated from the mature (processed) polypeptide sequence using the ProteinAnalysis module of Biopython (v1.87; Bio.SeqUtils.ProtParam), which computes average isotopic mass from amino acid composition; values are reported in kDa. For mitochondrial matrix proteins whose curated mature sequence was unavailable (54/218 Hsp60 interactome members), the full-length canonical sequence and its annotated transit-peptide/signal-peptide boundary were retrieved from UniProt (REST API, batch accession queries), and the mature sequence was obtained computationally by removing the N-terminal residues up to the annotated cleavage position prior to mass calculation, so that all mature-protein masses were derived consistently regardless of source.

*Reference proteome and interactome datasets*

The *E. coli* K-12 reference proteome (UniProt proteome ID UP000000625; reviewed/Swiss-Prot entries only, n = 4,403) was used as the genome-wide size background for GroEL substrates, with mass retrieved directly from the UniProt Mass field. The mitochondrial matrix background (n = 356) comprised proteins in a curated, deduplicated set of mature mitochondrial matrix sequences that were not present in the Hsp60 interactome (n = 218); the Hsp60 interactome itself (with the ones that were present in the matrix data), restricted to GroEL Class III (n = 84, a manually annotated obligate/high-affinity substrate class), and the full mitochondrial matrix set served as the four groups compared for size distribution.

(1) Size-distribution comparison across GroEL Class III, *E. coli* proteome, Hsp60 interactome, and mitochondrial matrix background

Size distributions were visualized as area-normalized frequency curves over 12 fixed-width bins spanning the pooled mass range of all groups being compared in a given panel, allowing direct shape comparison independent of the differing sample sizes (n = 84–4,403), and as notched box-and-whisker plots on a log10-transformed mass axis to accommodate the right-skewed, multi-order-of-magnitude range of protein masses without distortion from extreme values.

For each substrate group and its corresponding background, three complementary, non-parametric tests were applied, since mass distributions were non-normal (right-skewed, heavy-tailed) and violated assumptions required for parametric alternatives:

(2) Enrichment of the chaperonin interactors were calculated by dividing the frequency of the chaperonin interactors in each bin by the background number in the non-interacting set in the same bin.

(3) GroEL Class III vs. Hsp60 interactome: testing for a difference in distribution width (spread)

To determine whether GroEL Class III substrates and Hsp60 interactome members differ in the shape of their size distribution beyond any difference in central tendency, we applied a decomposition strategy rather than relying on a single omnibus statistic:

The two-sample KS test was first used to establish whether the overall distributions differ ($D = 0.195$, $p = 0.017$). A two-sided Mann-Whitney U test was then used to test whether this reflects a shift in median mass; this was not significant ($p = 0.203$; median 37.7 kDa for GroEL Class III vs. 40.9 kDa for Hsp60 interactome), indicating that the KS-detected difference is not attributable to a location shift. Levene's test for homogeneity of variance was then applied to test whether the two groups differ in dispersion specifically. Levene's test computes, for each observation, the absolute deviation from its group's median (the Brown-Forsythe modification, robust to non-normal/skewed data, in contrast to the classical mean-centered Levene's test or an F-test for equality of variances, both of which assume normality and are sensitive to outliers/skew — an inappropriate assumption for protein mass data), and then performs a one-way ANOVA on these absolute deviations across groups. A significant result indicates unequal variances (heteroscedasticity) between groups. This test returned a significant result (Levene statistic = 15.31, $p = 1.1 \times 10^{-4}$; SD = 14.2 kDa for GroEL Class III vs. 26.0 kDa for the Hsp60 interactome, $n = 84$ and 218 respectively).

***Calculation of the divergence of Hsp60 interactome***

*Protein Set Curation*

Mitochondrial matrix proteins were obtained from MitoCarta 3.0, filtering for proteins annotated to the matrix compartment. Canonical human protein sequences were retrieved from UniProtKB. Mitochondrial targeting sequences (MTS) were removed using a two-stage approach. First, experimentally annotated transit peptide cleavage sites were retrieved programmatically for each protein via the UniProt REST API. For proteins lacking an annotated transit peptide in UniProt, MTS cleavage was predicted computationally using TargetP 2.0 (Armenteros et al., 2019), submitted via the web server (services.healthtech.dtu.dk/services/TargetP-2.0/). Proteins predicted by TargetP 2.0 as non-mitochondrial were retained with their full sequence. The resulting mature sequences were deduplicated to remove redundant entries prior to downstream analysis.

*Ortholog Retrieval*

Strict one-to-one orthologs of each mature human matrix protein were retrieved across ten metazoan species representing major vertebrate and invertebrate lineages: Mus musculus, Rattus norvegicus, Canis lupus familiaris, Bos taurus, Monodelphis domestica, Gallus gallus, Xenopus tropicalis, Danio rerio, Drosophila melanogaster, and Caenorhabditis elegans. These species were selected to sample evolutionary distances spanning placental mammals, marsupials, birds, amphibians, teleost fish, and invertebrate ecdysozoans, thereby capturing a broad range of divergence timescales while remaining within Metazoa. Human UniProt accessions were first mapped to Ensembl gene identifiers via the UniProt ID Mapping API. Orthologous protein sequences were then retrieved using the Ensembl REST API (rest.ensembl.org), querying the homology endpoint for each human gene with type=orthologues, sequence=protein, and aligned=0, and retaining only relationships classified as ortholog_one2one. Proteins for which fewer than three orthologous sequences could be retrieved were excluded from further analysis.

*Multiple Sequence Alignment*

For each protein, the human mature sequence and its retrieved orthologs were aligned using MAFFT v7 in L-INS-i mode (--localpair --maxiterate 1000). The resulting multiple sequence alignment (MSA) was trimmed to the coordinates of the human reference sequence, removing alignment columns corresponding to insertions in non-human species. This anchoring procedure ensures that all positional entropy values are mapped to human residue coordinates.

*Positional Sequence Divergence*

Per-residue sequence divergence was quantified as Shannon entropy, calculated independently for each column of the human-anchored MSA. For a given alignment column, the observed amino acid frequency distribution P was computed across the non-gap characters of all sequences in the group, excluding alignment gaps and non-standard residues. Shannon entropy was calculated as $H = -\Sigma\ p_i \log_2(p\_i)$ over the

20 standard amino acids. Alignment columns corresponding to gaps in the human reference sequence were excluded, so entropy values are defined exclusively at human residue positions. Jensen-Shannon divergence relative to BLOSUM62 background amino acid frequencies was computed in parallel as an orthogonal measure of conservation. To reduce positional noise, raw per-residue Shannon entropy values were smoothed with a sliding window mean of 15 residues, applied symmetrically and centered on each position (minimum one observation required at boundaries). The smoothed values were stored per protein in a master residue-level table, with each row recording the UniProt identifier, absolute residue position, number of orthologs, and smoothed entropy.

*Length-Normalised Divergence Profiles and Statistical Comparison*

To compare divergence patterns across proteins of varying lengths, the absolute residue position of each entry was mapped to a fractional coordinate in [0, 1] using (position − 1) / (length − 1), where length is the number of residues in the mature human sequence. Matrix proteins were partitioned into two cohorts: those present in a curated Hsp60 interactome dataset (Hsp60 Interactome, ~270 proteins) and all remaining matrix proteins (Matrix Background). The fractional coordinate axis was discretised into 100 equal bins. For each bin, the distributions of smoothed Shannon entropy values from the Hsp60 Interactome and Matrix Background cohorts were compared using a two-sided Mann-Whitney U test. The resulting p-values were corrected for multiple comparisons using the Benjamini-Hochberg false discovery rate (FDR) procedure at $\alpha = 0.05$. To reduce spurious single-bin detections, only statistically significant bins (FDR < 0.05) forming a contiguous run of at least three consecutive bins were retained as significant regions. Mean disorder profiles per cohort were plotted with standard error of the mean as a function of normalised sequence position.

*Intrinsic Disorder Analysis*

As a parallel structural property, per-residue intrinsic disorder scores were predicted for each human mature matrix sequence using IUPred3 (Erdős et al., 2021) in long mode, which predicts long-range disorder. Predictions were computed locally using the IUPred3 standalone software. Disorder scores were smoothed with the same 15-residue sliding window applied to the divergence analysis, applied prior to length normalisation to maintain consistency. The smoothed disorder scores were subjected to the identical length-normalisation, binning, and statistical testing pipeline described above, enabling direct positional comparison between regions of differential disorder and regions of differential sequence divergence between the two cohorts.

***Chaperone cavity size estimation***

ATP bound structures of GroEL/ES or hHsp60/10 were obtained from Protein Data Bank (PDB IDs: 8BM1 and 9ES1) (85, 86). Structures were prepared by removing crystallographic water molecules and reformatting the PDB files for AMBER compatibility using *pdb4amber*. Topology files were generated with *tleap* using the AMBER ff19SB force field (87). Parameters for ATP were generated separately using *antechamber* and *parmchk2* modules within AmberTools, while $Mg^{2+}$ parameters were adopted from Allner *et al.* (88, 89). Systems were solvated with the TIP3P water model (90) and the resulting coordinates were used to calculate the cavity radius and volume using CICLOP (91, 92).

#### KEY RESOURCE TABLE

| REAGENT or RESOURCE | SOURCE | IDENTIFIER |
|---|---|---|
| **Antibodies** | | |
| Rabbit anti- GFP | Abcam | AB290 |
| Rabbit anti- MBP | NEB | E8030S |
| Rabbit anti- Hsp60 | CST | 4870S |
| Mouse anti-GAPDH | Elabsciences | E-AB-48016 |
| Anti- Rabbit IgG | CST | 7074S |
| **Bacterial and Viral strains** | | |
| *E.coli* DH5α | | |
| *E.coli* BL21 (DE3) Rosetta | | |
| *E.coli* WT (K-12, BW25113) (F-, *Δ(araD-araB)567*, *ΔlacZ4787*(::rrnB-3), *λ*-, *rph-1*, *Δ(rhaD-rhaB)568*, *hsdR514*) | CGSC (82) | CGSC#7636 |
| *E. coli ΔdnaK* (JW0013) (F-, *ΔdnaK734::kan, Δ(araD-araB)567, ΔlacZ4787(::rrnB-3), λ-, rph-1, Δ(rhaD-rhaB)568, hsdR514*) | CGSC (82) | CGSC#8342 |
| *E. coli ΔdnaJ* (JW0014) (F-, *ΔdnaJ735::kan*, *Δ(araD-araB)567*, *ΔlacZ4787*(::rrnB-3), *λ*-, *rph-1*, *Δ(rhaD-rhaB)568*, *hsdR514*) | CGSC (82) | CGSC#8343 |
| *E. coli Δtig* (JW0426) (F-, *Δ(araD-araB)567*, *ΔlacZ4787*(::rrnB-3), *Δtig-722::kan*, *λ*-, *rph-1*, *Δ(rhaD-rhaB)568*, *hsdR514*) | CGSC (82) | CGSC#8589 |
| *E. coli ΔhtpG* (JW0462) (F-, *Δ(araD-araB)567*, *ΔlacZ4787*(::rrnB-3), *ΔhtpG757::kan*, *λ*-, *rph-1*, *Δ(rhaD-rhaB)568*, *hsdR514*) | CGSC (82) | CGSC#8616 |
| **Chemicals, Peptides, and Recombinant Proteins** | | |
| NADH | Sigma | N8129 |
| ATP | Sigma | A2383 |
| Aluminium chloride | Sigma | 237078 |
| Sodium fluoride | Sigma | 201154 |
| Pyruvate Kinase/ Lactate Dehydrogenase | Sigma | P0294 |
| Phosphoenol pyruvate | Sigma | 860077 |
| SUPERase.In | Thermo Fisher Scientific | AM2696 |
| cOmplete His-Tag Purification Resin | Roche | 05893801001 |
| Butyl Sepharose 4 Fast Flow | Cytiva | 17098010 |
| DEAE Sepharose Fast Flow | Cytiva | 17070910 |
| Amylose resin | NEB | E8021S |
| HiLoad 16/600 Superdex 200 pg | Cytiva | 28989335 |
| Superose 6 Increase 10/300 GL | Cytiva | 29091596 |
| **Oligonucleotides** | | |

| | | |
|---|---|---|
| pBAD-SUMO-sGFP-mCherry-Insert_FP: TAGCAGGAGGAATTCGTCGACATGTCGGACTCAGAAGTC | This paper | KC 108 |
| pBAD-SUMO-sGFP-mCherry-Insert_RP: TTCACCTTTAGACATGCCCTGAAAATACAGGTTTTCGGTACC GGATCCACCAATCTGTTC | This paper | KC 109 |
| pBAD-sGFP-SUMO-mCherry-Insert_FP: GATGAATTGTACAAAGAAAACCTGTATTTTCAGGGCGTCGAC ATGTCGGACTCAGAAG | This paper | KC 113 |
| pBAD-sGFP-SUMO-mCherry-Insert_RP: GGGTATGGAAAGCTTGGTACCTTAGGATCCACCAATCTG | This paper | KC 114 |
| pBAD-sGFP-mCherry-fusion_FP: ATGGTGAGCAAGGGCGAGG | This paper | KC 116 |
| pBAD-sGFP-mCherry-fusion_RP: GCCCTTGCTCACCATGCCCTGAAAATACAGGTTTTCTTTGTAC AATTCATCCATACCATGG | This paper | KC 117 |
| pBAD-mCherry-sGFP-fused_FP: TATGAAGTCGACATGGTGAGCAAGGGCGAGGAGG | This paper | KC 175 |
| pBAD-mCherry-sGFP-fused_RP: AAGCTTGGTACCTTACTTGTACAGCTCGTCCATG | This paper | KC 176 |
| pBAD-2xsGFP-mCherry_FP: GAATTCGTCGACATGTCTAAAGGTGAAG | This paper | KC 118 |
| pBAD-2xsGFP-mCherry_RP: AAGCTTGGTACCTTATTTGTACAATTCATC | This paper | KC 119 |
| pBAD-sGFP-DMMBP_FP: ATACATGTCGACATGAAAATCGAAGAA | This paper | KC 128 |
| pBAD-sGFP-DMMBP_RP: GCTAGCGGTACCTTACTTGGTGATACG | This paper | KC 129 |
| pBAD-DMMBP-sGFP_RP: GCTAGCGGTACCCTTGGTGATACGAGT | This paper | KC 130 |
| pET-His-sGFP_FP: CGTCATATGGCTAGCATGTCTAAAGGTGAAGAATTATTCAC | This paper | KC 154 |
| pET-His-sGFP_RP: GCTAGCCATATGACGACCTTCGATATG | This paper | KC 155 |
| pET-sGFP-SUMO_FP: GGAATTCGCTAGCATGTCTAAAGGTGAAGAATTATTC | This paper | KC 156 |
| pET-sGFP-SUMO_RP: GGTACCAAGCTTTTAGGATCCACCAATCTGTTCTCTG | This paper | KC 157 |
| pET-DMMBP-sGFP_FP: CGTCGACGCTAGCATGAAAATCGAAGAAGGTAAAC | This paper | KC 158 |
| pET-DMMBP-sGFP_RP: TATGGAAAGCTTTTATTTGTACAATTCATCCATACC | This paper | KC 159 |
| pET-sGFP-DMMBP_FP: GAATTCGCTAGCATGTCTAAAGGTGAAGAATTATTC | This paper | KC 160 |
| pET-sGFP-DMMBP_RP: GGTACCAAGCTTTACTTGGTGATACGAGTCTGCG | This paper | KC 161 |
| GroEL_delES_FP: ATGGCAGCTAAAGACGTAAAATTC | This paper | KC 149 |
| GroEL_delES_RP: GTCTTTAGCTGCCATATGTATATCTCCTTCTTAAAGTTAAAC | This paper | KC 150 |
| hHsp60_delHsp10_FP: ATGTATGCCAAAGATGTGAAATTCGGTG | This paper | KC 151 |
| hHsp60_delHsp10_RP: ATCTTTGGCATACATATGTATATCTCCTTCTTAAAGTTAAAC | This paper | KC 152 |
| hHsp60_W42A_FP: GTTATTATTGAACAGAGCGCAGGTAGCCCGAAAGTG | This paper | SR 10 |

| | | |
|---|---|---|
| hHsp60_W42A_RP: CACTTTCGGGCTACCTGCGCTCTGTTCAATAATAAC | This paper | SR 11 |
| hHsp60_V72I_FP: CATTGGTGCCAAACTGATTCAGGATGTTGCCAATAAC | This paper | SR 14 |
| hHsp60_V72I_RP: GTTATTGGCAACATCCTGAATCAGTTTGGCACCAATG | This paper | SR 15 |
| hHsp60_Y201A_FP: GGTTATATTAGCCCGGCATTTATCAATACCAGC | This paper | SR 16 |
| hHsp60_Y201A_RP: GCTGGTATTGATAAATGCCGGGCTAATATAACC | This paper | SR 17 |
| hHsp60_S206A_FP: GTACTTTATCAATACCGCAAAAGGTCAGAAATGC | This paper | SR 18 |
| hHsp60_S602A_RP: GCATTTCTGACCTTTTGCGGTATTGATAAAGTAC | This paper | SR 19 |
| hHsp60_Y359A_FP: GTTACCACGAGCGAAGCAGAAAAAGAAAAACTG | This paper | SR 24 |
| hHsp60_Y359A_RP: CAGTTTTTCTTTTTCTGCTTCGCTCGTGGTAAC | This paper | SR 25 |
| pBAD-sGFP-Nterm _FP: TATGCCATAGCATTTTTATC | This paper | KC 124 |
| pBAD-sGFP-Nterm _RP: AGGTCAATTTACCGTAAGTAG | This paper | KC 125 |
| pBAD-sGFP-Cterm _FP: CAGACAACCATTACTTATCC | This paper | KC 126 |
| pBAD-sGFP-Cterm _RP: TTGGCGGTCTGGGTG | This paper | KC 127 |
| pETSmt3-sGFP-Nterm _FP: CCCCTCTAGAAATAATTTTG | This paper | KC 122 |
| pETSmt3-sGFP-Nterm _RP: TTCACCGGAGACAGAA | This paper | KC 123 |
| pETSmt3-sGFP-Cterm_RP: TTAGAGGCCCCAAGGGGTTATGCTA | This paper | KC 70 |
| pOFX-ES_FP: CCATAAACTGCCAGGAATTGGGG | This paper | KC 153 |
| ON-TARGETplus Human HSPD1 siRNA- SMART pool | Dharmacon | L-010600-00-0005 |
| **Recombinant DNA** | | |
| pBAD-yeGFP-mCherry | (29) | |
| pBAD-sGFP-mCherry | (29) | |
| pBAD-SUMO-sGFP-mCherry | This paper | |
| pBAD-sGFP-SUMO-mCherry | This paper | |
| pBAD-mCherry-sGFP-fused | This paper | |
| pBAD-sGFP-mCherry-fused | This paper | |
| pBAD-DMMBP-sGFP-mCherry | This paper | |
| pBAD-sGFP-DMMBP-mCherry | This paper | |
| pBAD-2x-sGFP-mCherry | This paper | |
| pBAD-NFsGFP-sGFP-mCherry | This paper | |
| pBAD-sGFP-NFsGFP-mCherry | This paper | |
| pBAD-MBP-sGFP-mCherry | This paper | |
| pBAD-sGFP-MBP-mCherry | This paper | |
| pET-His-sGFP | This paper | |
| pETSmt3-sGFP | (29) | |
| pET-SUMO-sGFP | This paper | |

| | | |
|---|---|---|
| pET-sGFP-SUMO | This paper | |
| pET-DMMBP-sGFP | This paper | |
| pET-sGFP-DMMBP | This paper | |
| pOFX-GroEL/ES | (29) | |
| pOFX-humanHsp60/10 | (30) | |
| pOFX-GroEL-delES | This paper | |
| pOFX-humanHsp60-delHsp10 | This paper | |
| pOFX-humanHsp60(W42A)/10 | This paper | |
| pOFX-humanHsp60(V72I)/10 | This paper | |
| pOFX-humanHsp60(Y201A)/10 | This paper | |
| pOFX-humanHsp60(S206A)/10 | This paper | |
| pOFX-humanHsp60(Y359A)/10 | This paper | |
| pOFX-aavEL/ES | This paper | |
| pOFX-aehEL/ES | This paper | |
| pOFX-aquHsp60/10 | This paper | |
| pOFX-athHsp60/10 | This paper | |
| pOFX-cflEL/ES | (30) | |
| pOFX-cmeHsp60/10 | This paper | |
| pOFX-crgHsp60/10 | This paper | |
| pOFX-ddiHsp60/10 | This paper | |
| pOFX-dmeHsp60/10 | This paper | |
| pOFX-glaHsp60/10 | This paper | |
| pOFX-hhaEL/ES | This paper | |
| pOFX-nveHsp60/10 | This paper | |
| pOFX-yeastHsp60/10 | (30) | |
| pOFX-smhEL/ES | (30) | |
| pOFX-spuHsp60/10 | This paper | |
| pOFX-swiEL/ES | This paper | |
| pOFX-tbrHsp60/10 | This paper | |
| pOFX-tetHsp60/10 | This paper | |
| pOFX-wbrEL/ES | This paper | |
| **Software and Algorithms** | | |
| ***Databases and Web Resources*** | | |
| MitoCarta 3.0 | (93) | https://www.broadinstitute.org/mitocarta |
| UniProtKB | (94) | https://www.uniprot.org |
| UniProt ID Mapping API | (94) | https://rest.uniprot.org/idmapping |
| Ensembl genome database | (95) | https://www.ensembl.org |
| Ensembl REST API | (96) | https://rest.ensembl.org |
| TargetP 2.0 web server | (97) | https://services.healthtech.dtu.dk/services/TargetP-2.0/ |
| ***Software and Algorithms*** | | |
| AlphaFold2 | (98) | |
| MAFFT v7 (L-INS-i mode) | (99) | https://mafft.cbrc.jp/alignment/software/ |

| | | |
|---|---|---|
| IUPred3 (standalone, long mode) | (100) | https://iupred.unimelb.edu.au/ |
| Python 3 | Python Software Foundation | https://www.python.org |
| NumPy | (101) | https://numpy.org |
| pandas | (102) | https://pandas.pydata.org |
| SciPy (mannwhitneyu) | (103) | https://scipy.org |
| statsmodels (multipletests, FDR-BH) | (104) | https://www.statsmodels.org |
| matplotlib | (105) | https://matplotlib.org |
| seaborn | (106) | https://seaborn.pydata.org |
| Custom divergence and disorder analysis scripts | This paper | N/A |
| ***Other*** | | |
| GroEL Interactome List | (15) | N/A |
| Hsp60 interactome list | (54) | N/A |

#### REFERENCES (SUPPLEMENTAL)


82. T. Baba *et al.*, Construction of Escherichia coli K-12 in-frame, single-gene knockout mutants: the Keio collection. *Molecular systems biology* **2**, MSB4100050 (2006).

83. D. Poso, A. R. Clarke, S. G. Burston, A kinetic analysis of the nucleotide-induced allosteric transitions in a single-ring mutant of GroEL. *Journal of molecular biology* **338**, 969-977 (2004).

84. J. Weaver *et al.*, GroEL actively stimulates folding of the endogenous substrate protein PepQ. *Nature communications* **8**, 15934 (2017).

85. I. Tascón *et al.*, Structural basis for ATP-driven double-ring assembly of the human mitochondrial Hsp60 chaperonin. *bioRxiv*, 2025.2010. 2004.680452 (2025).

86. S. Torino, M. Dhurandhar, A. Stroobants, R. Claessens, R. G. Efremov, Time-resolved cryo-EM using a combination of droplet microfluidics with on-demand jetting. *Nature methods* **20**, 1400-1408 (2023).

87. C. Tian *et al.*, ff19SB: amino-acid-specific protein backbone parameters trained against quantum mechanics energy surfaces in solution. *Journal of chemical theory and computation* **16**, 528-552 (2019).

88. O. Allnér, L. Nilsson, A. Villa, Magnesium ion–water coordination and exchange in biomolecular simulations. *Journal of chemical theory and computation* **8**, 1493-1502 (2012).

89. D. A. Case *et al.*, AmberTools. *Journal of chemical information and modeling* **63**, 6183-6191 (2023).

90. P. Mark, L. Nilsson, Structure and dynamics of the TIP3P, SPC, and SPC/E water models at 298 K. *The Journal of Physical Chemistry A* **105**, 9954-9960 (2001).

91. A. X. Francis *et al.*, An Algorithm-Based Investigation to Reveal the Differential Dynamics of Water inside Protein Cavities as a Function of Distance from Their Wall. *Journal of Chemical Information and Modeling* **65**, 5176-5187 (2025).

92. P. Garg, S. Sacher, P. Gautam, A. Ray, CICLOP: a robust and accurate computational framework for protein inner cavity detection. *Bioinformatics* **38**, 2153-2161 (2022).

93. S. Rath *et al.*, MitoCarta3. 0: an updated mitochondrial proteome now with sub-organelle localization and pathway annotations. *Nucleic acids research* **49**, D1541-D1547 (2021).

94. A. Bateman *et al.*, UniProt: the universal protein knowledgebase in 2023. *Nucleic Acids Res* **51**, D523-D531 (2023).

95. F. Cunningham *et al.*, Ensembl 2022. *Nucleic acids research* **50**, D988-D995 (2022).

96. A. Yates *et al.*, The Ensembl REST API: Ensembl data for any language. *Bioinformatics* **31**, 143-145 (2015).

97. J. J. Almagro Armenteros *et al.*, Detecting sequence signals in targeting peptides using deep learning. *Life science alliance* **2**, e201900429 (2019).

98. J. Jumper *et al.*, Highly accurate protein structure prediction with AlphaFold. *nature* **596**, 583-589 (2021).

99. K. Katoh, D. M. Standley, MAFFT multiple sequence alignment software version 7: improvements in performance and usability. *Molecular biology and evolution* **30**, 772-780 (2013).

100. G. Erdős, M. Pajkos, Z. Dosztányi, IUPred3: prediction of protein disorder enhanced with unambiguous experimental annotation and visualization of evolutionary conservation. *Nucleic acids research* **49**, W297-W303 (2021).

101. C. R. Harris *et al.*, Array programming with NumPy. *nature* **585**, 357-362 (2020).

102. W. McKinney, Data structures for statistical computing in Python. *scipy* **445**, 51-56 (2010).

103. P. Virtanen *et al.*, SciPy 1.0: fundamental algorithms for scientific computing in Python. *Nature methods* **17**, 261-272 (2020).

104. S. Seabold, J. Perktold, Statsmodels: econometric and statistical modeling with python. *scipy* **7**, 92-96 (2010).

105. J. D. Hunter, Matplotlib: A 2D graphics environment. *Computing in science & engineering* **9**, 90-95 (2007).

106. M. L. Waskom, Seaborn: statistical data visualization. *Journal of open source software* **6**, 3021 (2021).

#### SUPPLEMENTARY FIGURE LEGENDS

**Figure S1. Characterisation of GFP folding reporters and chaperonin activity. Supporting data for Figure 1.**

**(A)** Left, flow cytometry analysis of yeGFP and sGFP expressed in *E. coli* Wtk cells, demonstrating the reduced fluorescence of the slow-folding GFP (sGFP) reporter. Right, flow cytometry analysis of sGFP in the presence of GroEL/ES showing increased fluorescence upon chaperone expression. **(B)** Folding of sGFP expressed in the absence (spontaneous) or presence of GroEL/ES or hHsp60/10, measured by flow cytometry. Representative GFP, mCherry and GFP/mCherry distributions are shown. **(C)** Requirement of the cognate co-chaperonin (GroES or hHsp10) for folding assistance. **(D)** Cavity volume (left) and radius (right) plotted as a function of position along the longitudinal axis of the complex for hHsp60/10 and GroEL/ES cis ring. The ATP-bound hHsp60/10 chamber and the cis chamber of GroEL/ES exhibit similar volume and radius distributions, indicating comparable cavity dimensions.

**Figure S2. Characterisation of substrate encapsulation and folding. Supporting data for Figure 2.**

**(A)** Fluorescence anisotropy of free and GroEL/ES-encapsulated sGFP and SUMO-sGFP. **(B)** SDS-PAGE analysis of the solubility of MetK-yeGFP and yeGFP-MetK expressed in the absence (spontaneous) or presence of hHsp60/10. Soluble (S) and insoluble (P) fractions are shown.

**Figure S3. Expression and assembly of hHsp60 mutants. Supporting data for Figure 3.**

**(A)** SDS-PAGE analysis demonstrating comparable expression of wild-type and mutant hHsp60 together with hHsp10 in Wtk strain. **(B)** Native PAGE analysis of hHsp60/10 mutants following over-expression in Wtk cells, confirming assembly of oligomeric chaperonin complexes. Purified SR1 mutant of GroEL was included as a marker for heptameric or tetradecameric assemblies.

**Figure S4. Characterisation of chaperonin assembly, ATPase activity and folding assays. Supporting data for Figure 4.**

**(A)** Native PAGE analysis of GroEL/ES and hHsp60/10 following over-expression in Wtk cells, confirming assembly of oligomeric chaperonin complexes. Purified SR1 mutant of GroEL was included as a marker for heptameric or tetradecameric assemblies. **(B)** ATP hydrolysis rates of purified GroEL/ES and hHsp60/10, expressed per μM of tetradecameric chaperonin. **(C)** Apparent refolding rates of sGFP and SUMO-sGFP following dilution from denaturant in the presence of clarified whole-cell lysates from cells over-expressing GroEL/ES or hHsp60/10. Neither lysate recapitulates the preferential assistance of SUMO-sGFP observed *in vivo*. **(D)** Folding of reciprocal constructs with MBP and DMMBP fused to sGFP by hHsp60/10. **(E)** Immunoblots of input (I) and amylose-bound (B) fractions following *in vitro* translation of sGFP-DMMBP and DMMBP-sGFP in the absence (spontaneous) or presence of hHsp60/10. Folded DMMBP was pulled-down by amylose affinity chromatography and detected using an anti-MBP antibody. Data are mean ± SD from three independent experiments. Statistical significance was determined using two-tailed Student's t-tests; p-values depicted on top.

**Figure S5. Characterisation of mitochondrial folding reporters and sequence properties of hHsp60 interactors. Supporting data for Figure 5.**

**(A)** Western blot confirming siRNA mediated knockdown of Hsp60 in HeLa cells. GAPDH served as the loading control. **(B)** Representative confocal images of HeLa cells expressing mitochondrially targeted DMMBP-sGFP or sGFP-DMMBP following control or HSPD1 (Hsp60) knock down. Right, Quantification of mean GFP fluorescence. Data are mean ± SD. **(C)** Average predicted disorder from sequence plotted along the normalised length of non-substrate matrix proteins (grey) and the hHsp60 interactome (red). Mitochondrial targeting sequences were excluded before analysis. Red-shaded regions indicate sequence segments showing significant differences in sequence divergence between the two protein sets (these are the same regions corresponding to the shaded regions in Figure 4A). Blue-shaded regions indicate sequence segments showing significant differences in predicted disorder between the two protein sets. **(D)** Number of unique proteins (left) and total number of residues in each length bin (right) considered for non-substrate matrix proteins (grey line) and Hsp60 interactome (red line) plotted against the normalized length of the protein. Shaded regions indicate sequence segments showing significant differences in sequence divergence between the two groups. The sequences were normalized between zero and one, after removing mitochondrial signal sequences.

**Figure S6. Expression and folding activity of chaperonin homologs. Supporting data for Figure 6.**

**(A)** SDS-PAGE analysis demonstrating expression of GroEL/Hsp60 homologs together with their cognate GroES/Hsp10 co-chaperonins in Wtk strain. Homologs for which no detectable over-expression was observed are labelled in red. **(B)** *in vivo* folding assistance by all selected prokaryotic homologs. **(C)** *in vivo* folding assistance by all selected eukaryotic homologs. Data are mean ± SD from three independent experiments.

**Figure S1**

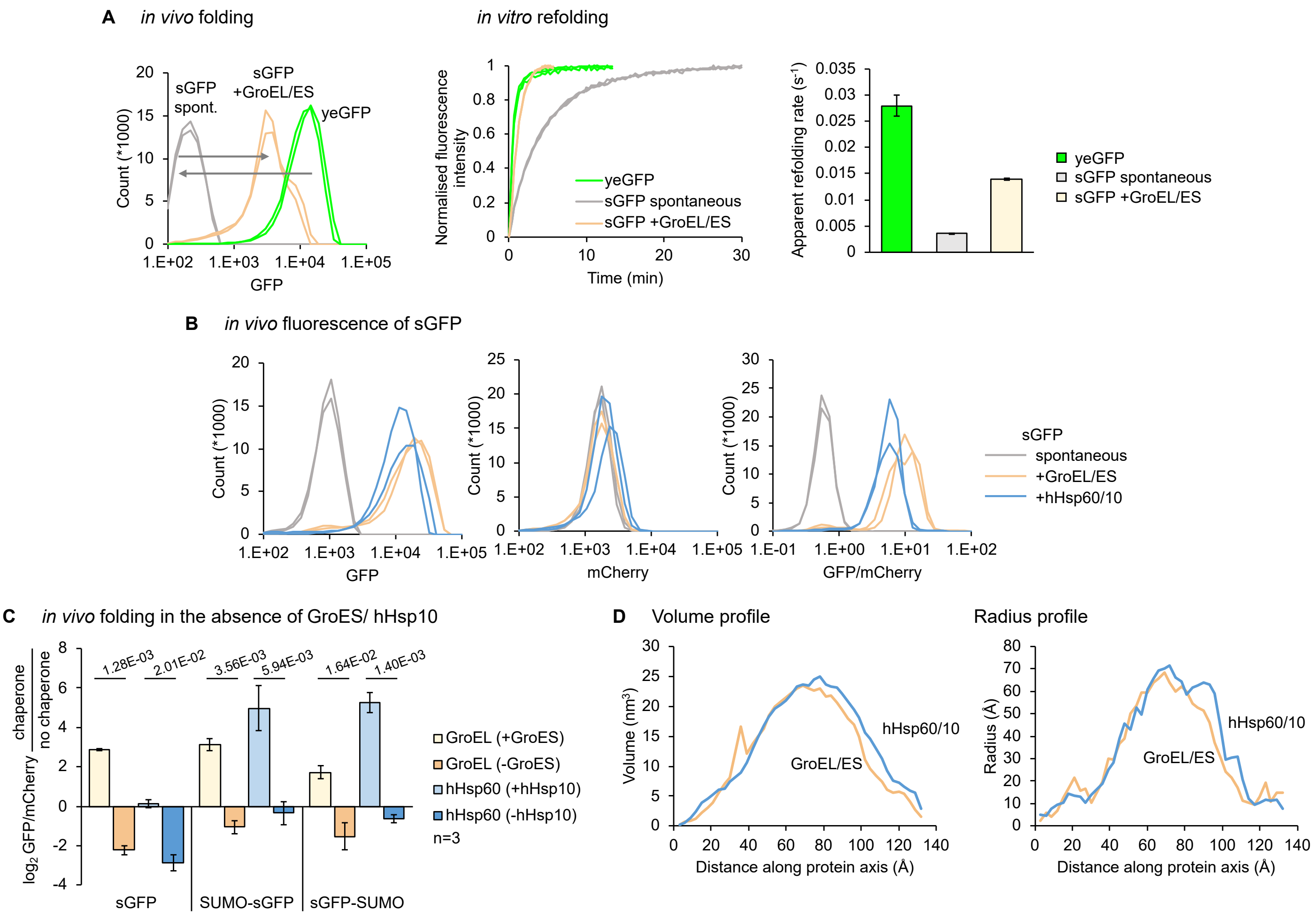
A
in vivo folding
in vitro refolding
sGFP spont.
sGFP +GroEL/ES
yeGFP
Count (*1000)
GFP
Normalised fluorescence intensity
Time (min)
yeGFP
sGFP spontaneous
sGFP +GroEL/ES
Apparent refolding rate (s-1)
B
in vivo fluorescence of sGFP
mCherry
GFP/mCherry
sGFP
spontaneous
+GroEL/ES
+hHsp60/10
C
in vivo folding in the absence of GroES/ hHsp10
1.28E-03
2.01E-02
3.56E-03
5.94E-03
1.64E-02
1.40E-03
GroEL (+GroES)
GroEL (-GroES)
hHsp60 (+hHsp10)
hHsp60 (-hHsp10)
n=3
sGFP
SUMO-sGFP
sGFP-SUMO
D
Volume profile
Radius profile
Volume (nm3)
Radius (Å)
hHsp60/10
GroEL/ES
Distance along protein axis (Å)

### Figure S2

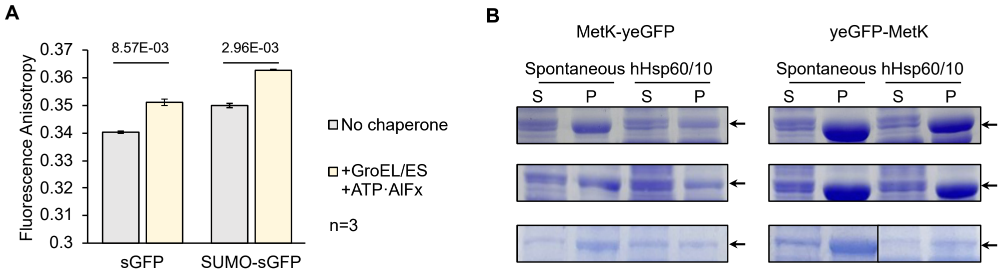
A
8.57E-03
2.96E-03
Fluorescence Anisotropy
0.37
0.36
0.35
0.34
0.33
0.32
0.31
0.3
sGFP
SUMO-sGFP
No chaperone
+GroEL/ES +ATP·AlFx
n=3
B
MetK-yeGFP
yeGFP-MetK
Spontaneous
hHsp60/10
S
P

**Figure S3**

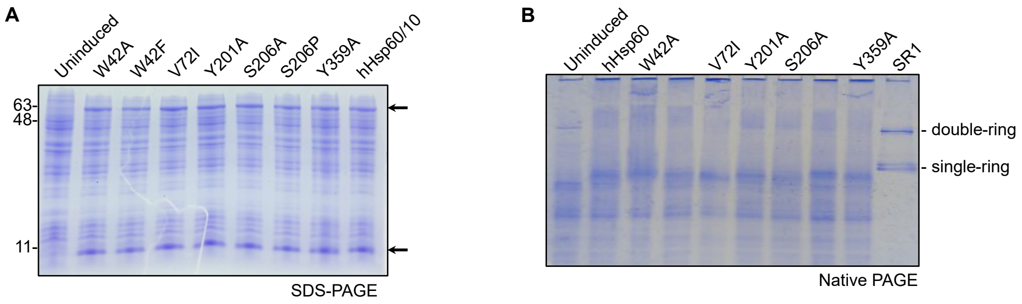
A
Uninduced
W42A
W42F
V72I
Y201A
S206A
S206P
Y359A
hHsp60/10
63-
48-
11-
SDS-PAGE
B
Uninduced
hHsp60
W42A
V72I
Y201A
S206A
Y359A
SR1
- double-ring
- single-ring
Native PAGE

#### Figure S4

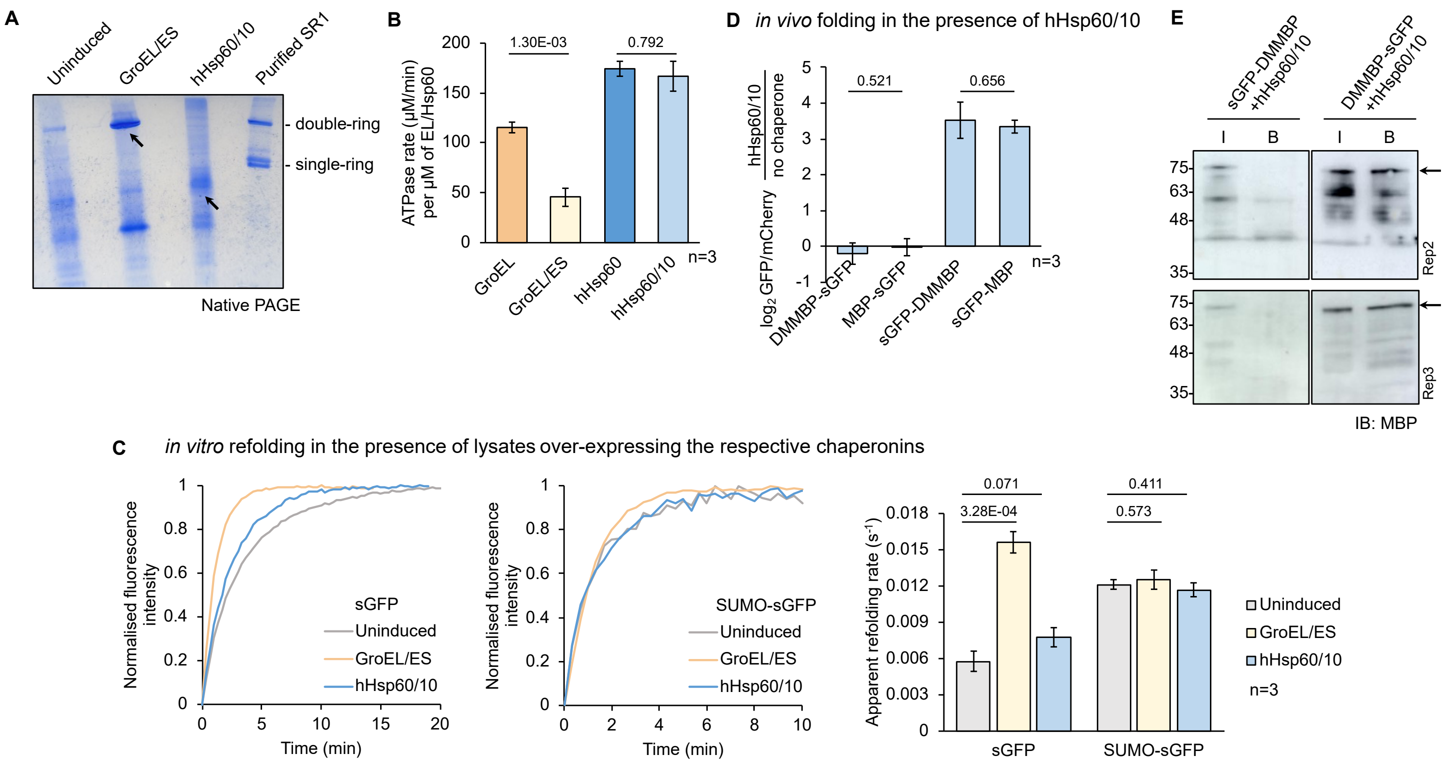
A
Uninduced
GroEL/ES
hHsp60/10
Purified SR1
- double-ring
- single-ring
Native PAGE
B
ATPase rate (μM/min) per μM of EL/Hsp60
1.30E-03
0.792
GroEL
GroEL/ES
hHsp60
hHsp60/10
n=3
D in vivo folding in the presence of hHsp60/10
log2 GFP/mCherry hHsp60/10 / no chaperone
0.521
0.656
DMMBP-sGFP
MBP-sGFP
sGFP-DMMBP
sGFP-MBP
n=3
E
sGFP-DMMBP +hHsp60/10
DMMBP-sGFP +hHsp60/10
I
B
Rep2
Rep3
IB: MBP
C in vitro refolding in the presence of lysates over-expressing the respective chaperonins
Normalised fluorescence intensity
Time (min)
sGFP
SUMO-sGFP
Uninduced
GroEL/ES
hHsp60/10
Apparent refolding rate (s-1)
0.071
3.28E-04
0.411
0.573
n=3

#### Figure S5

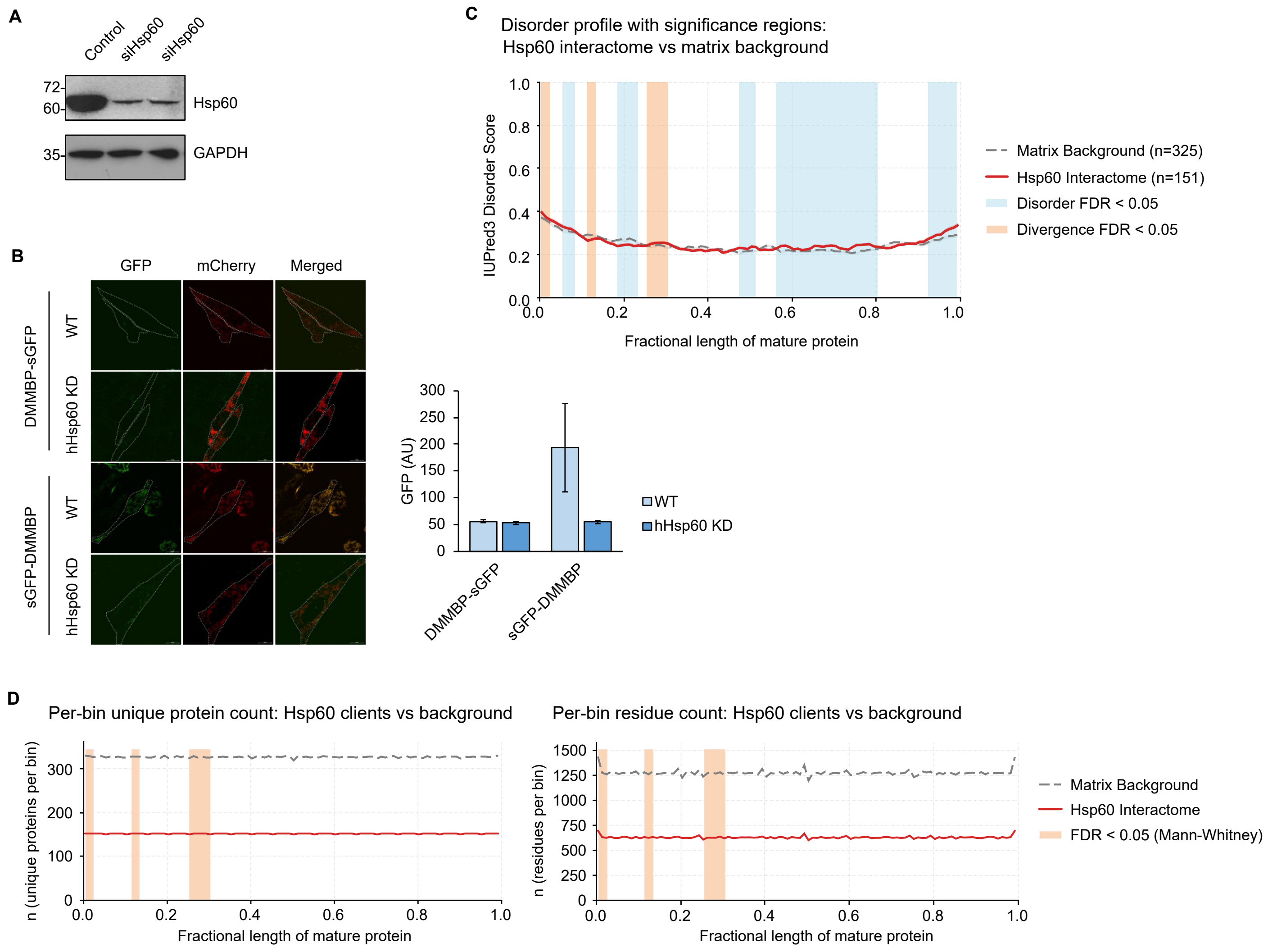

A
Control
siHsp60
siHsp60
72-
60-
Hsp60
35-
GAPDH
C
Disorder profile with significance regions:
Hsp60 interactome vs matrix background
IUPred3 Disorder Score
Fractional length of mature protein
Matrix Background (n=325)
Hsp60 Interactome (n=151)
Disorder FDR < 0.05
Divergence FDR < 0.05
B
GFP
mCherry
Merged
DMMBP-sGFP
sGFP-DMMBP
WT
hHsp60 KD
GFP (AU)
WT
hHsp60 KD
DMMBP-sGFP
sGFP-DMMBP
D
Per-bin unique protein count: Hsp60 clients vs background
n (unique proteins per bin)
Fractional length of mature protein
Per-bin residue count: Hsp60 clients vs background
n (residues per bin)
Fractional length of mature protein
Matrix Background
Hsp60 Interactome
FDR < 0.05 (Mann-Whitney)

**Figure S6**

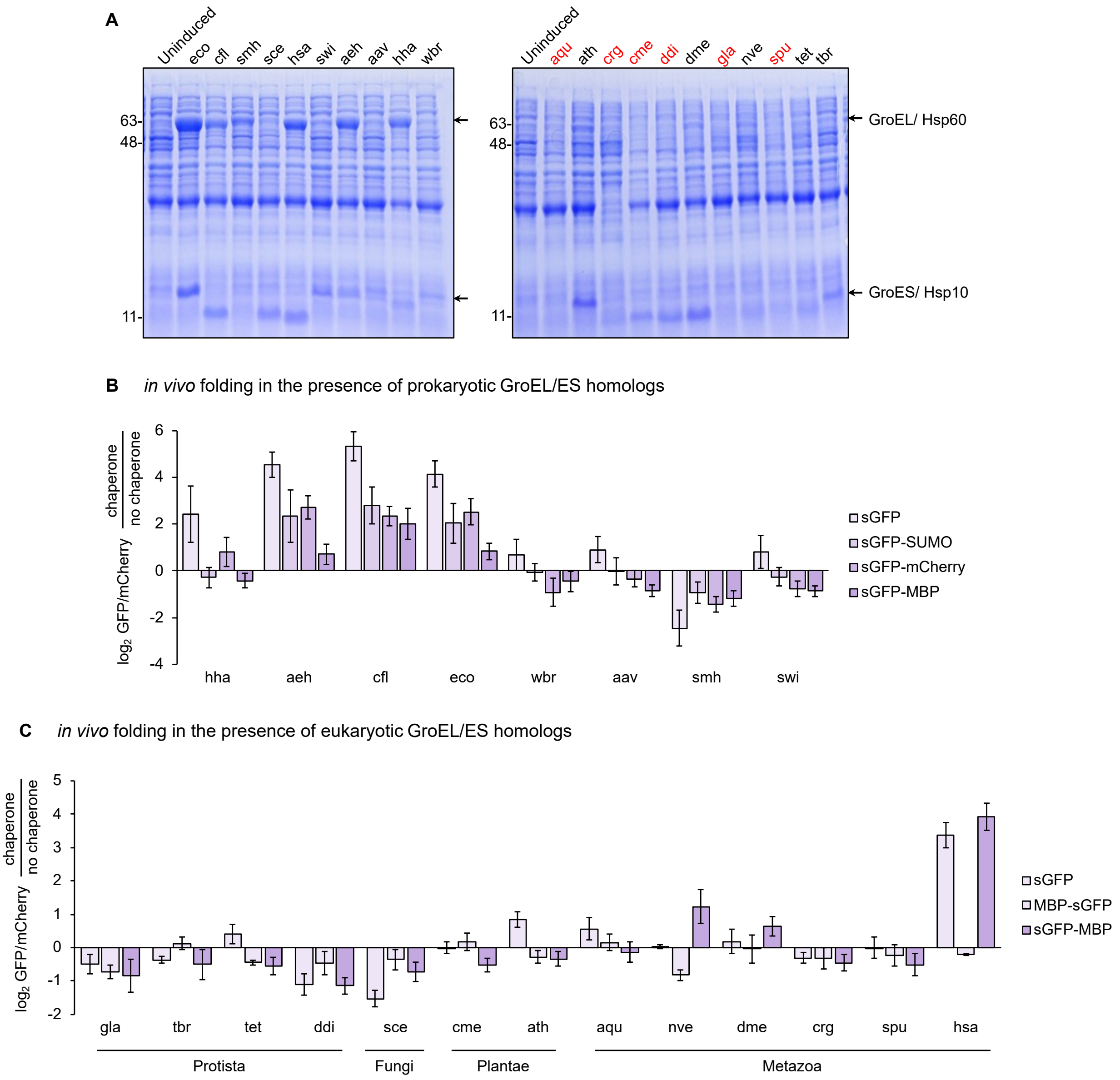
A
Uninduced
eco
cfl
smh
sce
hsa
swi
aeh
aav
hha
wbr
63-
48-
11-
Uninduced
aqu
ath
crg
cme
ddi
dme
gla
nve
spu
tet
tbr
63-
48-
11-
GroEL/ Hsp60
GroES/ Hsp10
B in vivo folding in the presence of prokaryotic GroEL/ES homologs
log2 GFP/mCherry chaperone/no chaperone
6
4
2
0
-2
-4
hha
aeh
cfl
eco
wbr
aav
smh
swi
sGFP
sGFP-SUMO
sGFP-mCherry
sGFP-MBP
C in vivo folding in the presence of eukaryotic GroEL/ES homologs
log2 GFP/mCherry chaperone/no chaperone
5
4
3
2
1
0
-1
-2
gla
tbr
tet
ddi
sce
cme
ath
aqu
nve
dme
crg
spu
hsa
Protista
Fungi
Plantae
Metazoa
sGFP
MBP-sGFP
sGFP-MBP
